# Direct readout of neural stem cell transgenesis with an integration-coupled gene expression switch

**DOI:** 10.1101/834424

**Authors:** Takuma Kumamoto, Franck Maurinot, Raphaëlle Barry, Célia Vaslin, Sandrine Vandormael-Pournin, Mickaël Le, Marion Lerat, Michel Cohen-Tannoudji, Alexandra Rebsam, Karine Loulier, Stéphane Nédelec, Samuel Tozer, Jean Livet

## Abstract

**SUMMARY:** Stable genomic integration of exogenous transgenes is critical for neurodevelopmental and neural stem cell studies. Despite the emergence of tools driving genomic insertion at high rates with DNA vectors, transgenesis procedures remain fundamentally hindered by the impossibility to distinguish integrated transgenes from residual episomes. Here, we introduce a novel genetic switch termed iOn that triggers gene expression upon insertion in the host genome, enabling simple, rapid and faithful identification of integration events following transfection with naked plasmids accepting large cargoes. In vitro, iOn permits rapid drug-free stable transgenesis of mouse and human pluripotent stem cells with multiple vectors. In vivo, we demonstrate accurate cell lineage tracing, assessment of regulatory elements and mosaic analysis of gene function in somatic transgenesis experiments that reveal new aspects of neural progenitor potentialities and interactions. These results establish iOn as an efficient and widely applicable strategy to report transgenesis and accelerate genetic engineering in cultured systems and model organisms.

## INTRODUCTION

Gene transfer approaches enabling stable genomic integration and expression of exogenous transgenes in host cells or organisms are central in biology. Constant progress in integrative vector systems facilitate their implementation for a growing range of purposes, such as investigation of gene function and regulation (Akhtar et al., 2013), genetic screens (Doench, 2018), stem cell engineering (Tewary et al., 2018), cell based therapies (Hirsch et al., 2017) and emerging synthetic biology applications (Black et al., 2017; Ebrahimkhani and Ebisuya, 2019).

Neurobiology is one of the fields which has been the most profoundly impacted by these approaches. In cultured systems, additive transgenesis is widely used to derive cells homogenously expressing one or more genes of interest, a process instrumental in efforts to direct pluripotent stem cells towards specific neuronal lineages, decipher the mechanisms regulating their differentiation and harness these cells to model neural pathologies (e.g. Kondo et al., 2017; Nehme et al., 2018; Yang et al., 2017). In vivo, somatic transgenesis approaches that directly target neural progenitors have acquired major importance for neurodevelopmental studies by enabling to permanently mark these cells and trace their lineage. Seminal studies have taken advantage of retroviral vectors to characterize the neuronal and glial descent of embryonic neural progenitors in model animals (Noctor et al., 2001; Price et al., 1987; Yu et al., 2012). The location of these progenitors along the lumen of the neural tube also makes them accessible to integrative schemes based on DNA electroporation applicable to trace neural cell lineage (Chen and Loturco, 2012; Loulier et al., 2014), target specific genes and mark their product (Mikuni et al., 2016; Suzuki et al., 2016), or even perform in situ screens for genes involved in neurodevelopmental processes (Lu et al., 2018). Postmitotic neurons can also be directly targeted with lentiviral vectors (Parr-Brownlie et al., 2015) or recent AAV-based strategies (Nishiyama et al., 2017).

Genomic integration at the highest possible rate is essential in all the above applications. Viral vectors are still broadly used to this effect, but their production is time and resource consuming and they are inherently limited in cargo capacity. Fast and convenient to produce, naked DNA vectors have become a tool of choice for gene transfer, combined with transposases or programmable endonucleases (Ivics et al., 2009; Suzuki et al., 2016), the former ones providing higher efficiency essential for the integration of large transgenes. However, stable transgenesis with classic DNA vectors faces a universal problem: the identification of genome-integration events after transfection is hampered by the presence of residual non-integrated (episomal) transgene copies. In cultured cells, week-long delays are required to eliminate these episomes, usually in presence of drugs, with inherent risks of genetic or epigenetic drift (Liang and Zhang, 2013; Merkle et al., 2017), severely limiting the applicability and throughput of stable transgenesis compared to transient expression approaches. In vivo, transient episomal expression may confound the interpretation of somatic transgenesis experiments, because the past history of transgene expression in a population of cells may not be accurately reflected by the markers they harbor: as episomal transgenes are progressively lost through cell division, they may alter the behavior of progenitors while being undetectable in their descent. This essentially precludes the development of reliable functional mosaic analysis schemes based on exogenous DNA vectors applicable to probe neural stem cell regulation, until now only possible by resorting to complex genetic schemes (Pontes-Quero et al., 2017; Zong et al., 2005). Moreover, in vitro as well as in vivo, episomes may cause leakage from cell type-specific promoters (Inoue et al., 2017) and the burst of marker expression that follows transfection may have harmful effects on cell behavior, identity and viability (Batard et al., 2001).

Here, we introduce an *<u>i</u>*ntegration-coupled *<u>On</u>* (iOn) gene expression switch through which genomic insertion of an initially silent transgene triggers its expression. We present several implementations of this concept that efficiently couple transcriptional or translational gene activation to genomic integration. We demonstrate its advantages in vitro for highly efficient establishment of stable mouse and human pluripotent stem cell lines expressing multiple transgenes. In vivo, somatic transfection with iOn is a powerful alternative to additive transgenesis to drive constitutive or conditional expression of reporters and effectors. This enabled us to determine the clonal output of progenitor subtypes in the developing retina and to uncover a homeostatic control of neurogenesis in the embryonic neural tube. These results establish iOn as an efficient strategy for direct readout of transgenesis applicable to accelerate and facilitate genetic manipulations in neurodevelopmental and neural stem cell studies.

## RESULTS

### Design and validation of an integration-coupled gene expression switch

To create the iOn switch (Figure 1), we took advantage of the piggyBac transposition system (Fraser et al., 1996), currently one of the most efficient tool for genomic integration of exogenous DNA (Ding et al., 2005), characterized by its very large cargo capacity (Li et al., 2011) and precise cut-and-paste mechanism mediating traceless transposon excision. In classic transposon vectors (Figure 1A, left), the transgene, framed by two antiparallel-oriented terminal repeats recognized by the piggyBac transposase, is readily active prior to excision from the donor plasmid. We reasoned that placing the terminal repeats in parallel rather than antiparallel orientation would create a situation in which transposase-mediated insertion in the host genome is accompanied by a rearrangement exploitable to trigger gene expression (Figure 1A, right).

**Figure 1.**
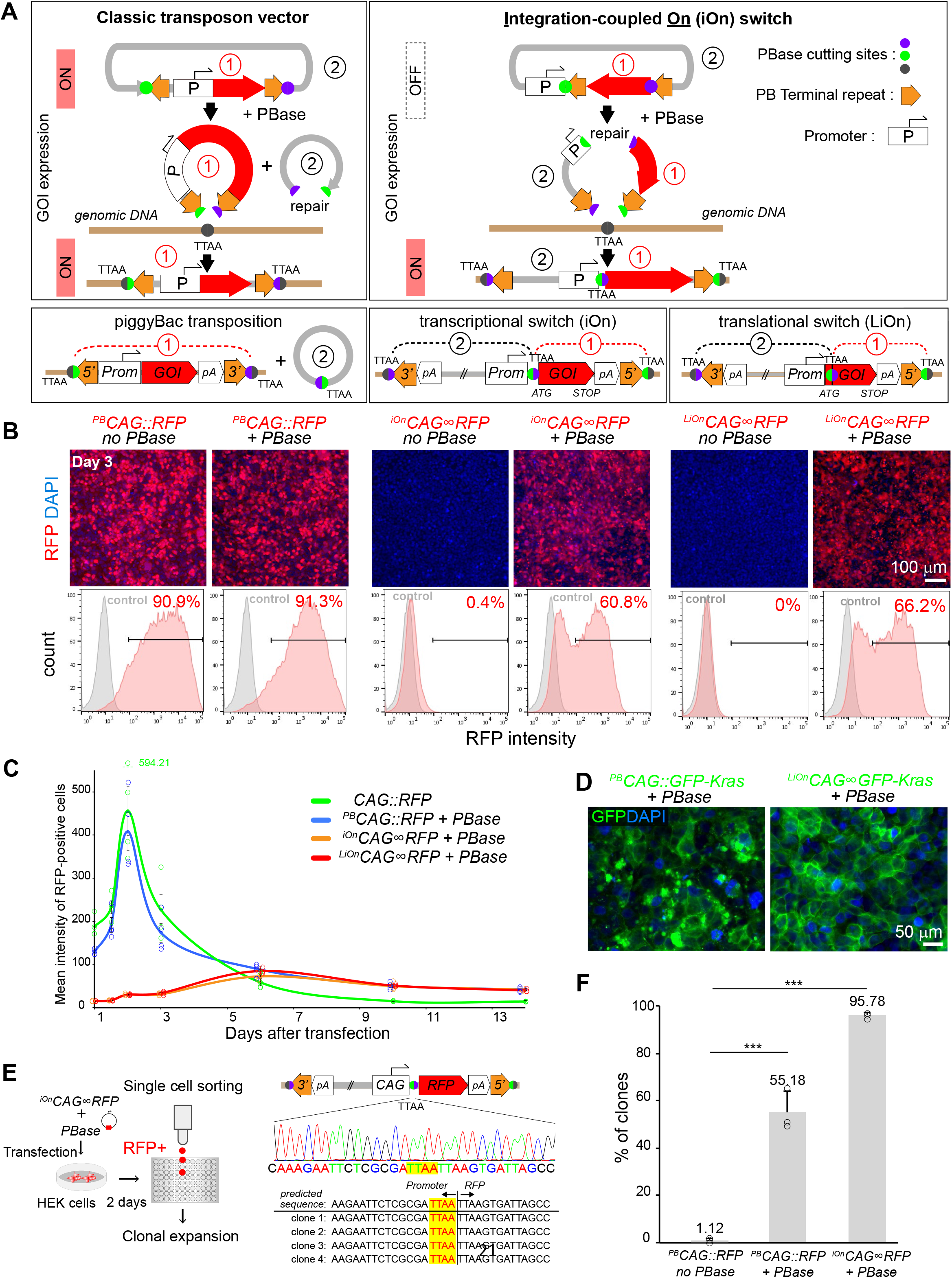
Principle and validation of the iOn switch. (A) Principle of gene transfer with classic transposon (left) and iOn vectors (right). While the former allow GOI expression from episomes, that from iOn vector is conditioned by transposase action which reunites either the promoter (Prom) and GOI (transcriptional switch) or 5’ and 3’ portions of the GOI (translational “LiOn” switch). 5’ and 3’: transposon terminal repeats; pA: transcription terminators; TTAA: PB transposition footprint. (B) Validation in HEK293 cells 3 days after transfection with a classic transposon (^*PB*^*CAG∷RFP*, left) and iOn (middle) or LiOn vectors (right). Top: epifluorescence images. Bottom: representative cytometry plots from cells transfected with PB/iOn vectors (red) vs. control cells (grey). (C) Time-course analysis of RFP expression with episomal, classic transposon and iOn/LiOn vectors. Values and error bars represent the mean and s.e.m. of four replicates. (D) Localization of a membrane-GFP (GFP-Kras) expressed from a classic transposon (left) or LiOn vector (right) 3 days after transfection in HEK293 cells. (E) Clones established by sorting ^*iOn*^*CAG∞RFP*-transfected cells display the sequence expected for precise junction between promoter and GOI. (F) Cells sorted based on ^*iOn*^*CAG∞RFP* expression yield a high proportion of RFP-positive clones compared to ^*PB*^*CAG∷RFP*. Values and error bars represent the mean and s.e.m. of 3 separate experiments. 1078, 620 and 504 clones were assessed for ^*PB*^*CAG∷RFP* transfection without and with PBase and ^*iOn*^*CAG∞RFP*, respectively (p<0.0001 with χ^2^ test). See also Figures S1 and S2.

We first designed transcriptional versions of iOn in which a promoter (Prom) and gene of interest (GOI), initially placed in opposite orientation, are reunited in a functional configuration by transposition (Figures 1A, middle, and S1). We denote this configuration ^*iOn*^*Prom∞GOI,* by opposition to a classic transposon driving constitutive expression noted ^*PB*^*Prom∷GOI*. We tested different iOn vector designs using the broadly active *CAG* promoter and a red fluorescent protein (RFP) as GOI, which we assayed by transfection in HEK293 cells in presence or absence of piggyBac transposase (PBase) (Figure S1). 3 days after transfection, PBase-dependent RFP expression was observed with all tested constructs, validating the iOn switch concept. We selected the configuration with highest signal to noise ratio (Figure S1C). This ^*iOn*^*CAG∞RFP* construct yielded efficient RFP expression with minimal background transcription in absence of PBase (0.4%, Figure 1B, middle). While this faint leakiness may be tolerated in most applications, we nevertheless sought to suppress it by designing a “Leak-proof iOn” (LiOn) switch in which both transcription and translation are blocked in absence of transposase. In LiOn vectors, the GOI open reading frame (ORF) is initially interrupted after the translational start and reconstituted upon PBase action, the piggyBac excision footprint (TTAA) being incorporated at a silent or neutral position (Figure 1A, bottom right). A vector based on this design (hereafter denoted ^*LiOn*^*Prom∞GOI*) showed undetectable leakiness in absence of transposase (Figure 1B, right). Time-course experiments comparing the two types of iOn vectors with classic piggyBac-based and non-integrative plasmids showed that the iOn and LiOn switches achieve long-term GOI expression without the transient expression burst associated with vectors active in episomal form (Figure 1C). Instead, GOI expression gradually builds up, plateaus within a few days and remains in a narrower range compared to a classic transposon (Figure 1C and Figure S2A,B), a feature of high interest to reduce variability and improve protein localization in transfection assays. Indeed in HEK293 cells transfected with a LiOn vector expressing a farnesylated GFP (GFP-Kras), near-uniform membrane expression was observed three days after transfection, while signal from a classic transposon showed frequent clustering and overflowing of GFP from the membrane compartment (Figure 1D). To confirm that the switch drives the expected transgene rearrangement, we derived clones from ^*iOn*^*CAG∞RFP*-expressing cells in which we sequenced the junction between the promoter and GOI. All sequences demonstrated reunion of the two transgene elements with base pair-precision (Figure 1E, n=6). Moreover, analysis of >500 clones showed that 95.78% (±0.73 SEM) remained RFP-positive 10 days after sorting, indicating highly efficient integration of the transgene in the genome of founder cells and maintenance of its expression over the long term (Figure 1F). By contrast, a classic ^*iOn*^*CAG∷RFP* vector yielded only 55.18% (±5.07 SEM) of RFP-positive clones. Similarly, in mouse ES cells, fluorescence-based clonal selection with the above ^*LiOn*^*CAG∞GFP-Kras* vector demonstrated a high enrichment in integrative events compared to a classic transposon (Figure S2C). Thus, the dependence of the iOn switch on transposition enables efficient coupling of transgene activation and genomic integration. Importantly, short and long-term viability with iOn vectors were comparable to that of classic transposons (Figure S2D,E), and the switch was active in all cells tested, including HeLa and 3T3 (Figure S2F) as well as human pluripotent stem cells (see below). These results establish iOn as a tool for highly efficient drug-free transgenesis through which genomic expression of GOIs can be assessed directly after transfection without interfering episomal expression.

### High efficiency multiplexed stable transfection with iOn vectors

Having validated the iOn switch concept, we sought to further test and extend its range of applications in cultured cells (Figure 2). Stable transfection with multiple independent transgenes is challenging with standard DNA vectors due to false positives resulting from episomal expression and the difficulty of combining orthogonal drug selection systems. The high enrichment for integrative events achieved with iOn bypasses these issues. To simultaneously identify multiple integration events, we created LiOn vectors expressing green and near-infrared fluorescent proteins (FPs), forming a trichromatic set together with the abovementioned ^*LiOn*^*CAG∞RFP* (Figure 2A and Figure S3A). Co-transfection of the three vectors in HEK293 cells resulted in varied FP combinations, revealing activity of multiple transgenes in a large fraction of cells (Figure 2B). Strikingly, sorting and expansion of triple-fluorescent cells yielded a vast majority (80%) of clones co-expressing all three FPs 10 days after sorting, compared to only about 20% when using classic transposons (Figure 2C and Figure S3B,C), demonstrating the superior efficiency of iOn-based multiplexed transgenesis. In human induced pluripotent stem (iPS) cells, clones co-expressing all three ^*LiOn*^*CAG∞FP* vectors similarly maintained their expression at near homogenous levels over multiple passages (Figure 2D). The three-color LiOn transgenes also drove long-term expression during iPS cells differentiation towards the neural lineage, with FP combinations providing a readout of clonal relationships (Figure 2E). Thus, iOn provides an efficient route for one-shot multiplexed transgenesis in cell culture models.

**Figure 2.**
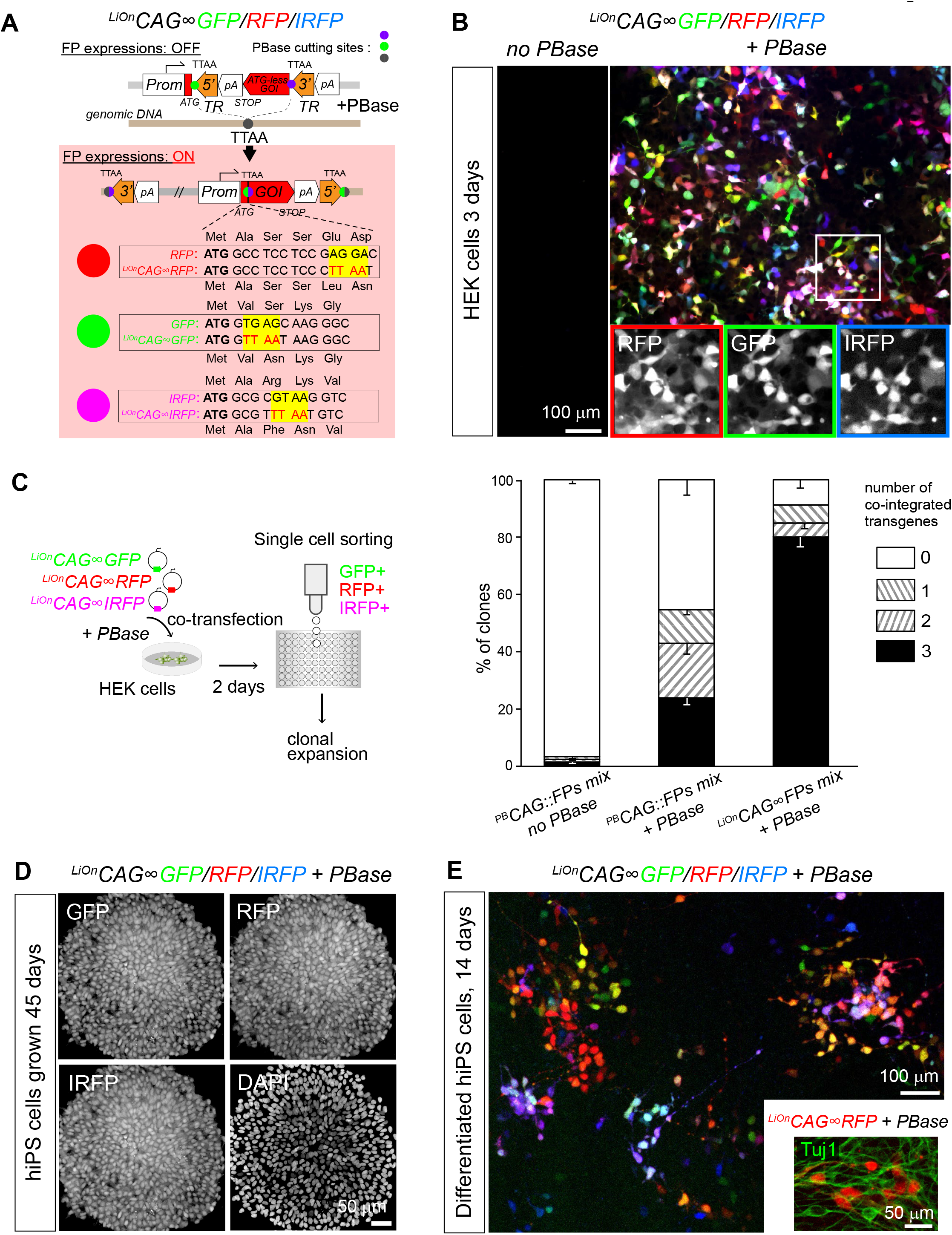
Highly efficient multiplexed stable transfection with iOn vectors. (A) Sequence design of LiOn plasmids expressing three distinct FPs: mRFP1, EGFP or IRFP670. (B) 3 days after co-transfection of the three ^*LiOn*^*CAG∞FP* plasmids in HEK293 cells, PBase-dependent expression is observed at similar levels for all FPs. (C) Cell sorting of triple-labeled cells 2 days after transfection with the LiOn vectors yields a majority of clones co-expressing the three FPs, but only a minority with classic transposons (mean and s.e.m. of three separate experiments; 158, 104 and 87 clones were assessed for each condition). χ^2^ test indicated significant difference between the three situations (p<0.0001). (D) Example of human iPS cell colony derived from cells co-transfected with the three-color ^*LiOn*^*CAG∞FP* vectors, grown 45 days. All cells co-express the three FPs. (E) Co-transfection of the three-color ^*LiOn*^*CAG∞FP* vectors during human iPS cell neuronal differentiation yields varied FP combinations reflecting their clonal organization. Inset: ^*LiOn*^*CAG∞RFP* expression in iPS cells immunostained with the neuronal marker Tuj1 (green). See also Figures S3.

### Additive somatic transgenesis and cell lineage tracing with iOn

We next assessed the iOn switch in vivo (Figure 3). By efficiently coupling transgene integration and expression, iOn vectors create a situation akin to additive transgenesis through simple transfection. We tested the applicability of this strategy for somatic cell transgenesis in various assays in the developing vertebrate nervous system, where embryonic electroporation provides access to neural progenitors. In the mouse cerebral cortex, electroporation of a classic episomal vector (*CAG∷GFP*) only labeled neurons born at the time of the electroporation (E12) that migrated in intermediate layers, due to its rapid dilution in dividing progenitors (Figure 3A and S4A). In striking contrast, an ^*iOn*^*CAG∞RFP* vector homogenously marked electroporated progenitors and all their radially-migrating derivatives, including late born upper-layer neurons and astrocytes, in a strict PBase-dependent manner. Further tests in the embryonic chicken spinal cord confirmed that iOn expression in progenitor cells, first detected one day after electroporation, was maintained in their descent (Figure 3B and S4B,C). Importantly, it avoided the strong and irregular labeling of isolated neurons polluting patterns obtained with classic transposons and episomes, likely due to inheritance of multiple non-integrated copies in cells born shortly after transfection (Figure 3B). Similar to the above two models, iOn vector electroporation in the mouse retina at the start of neurogenesis successfully labeled all retinal layers, while episomes only marked early born retinal ganglion cells (Figure 3C). In addition, combining iOn vectors expressing different colors enabled to further resolve the lineage relationships of labeled cells (Figure S4D). In the chick retina, retinal pigmented epithelium and spinal cord, FP combinations generated by the trichromatic ^*LiOn*^*CAG∞FP* vectors efficiently contrasted groups of cells that matched known clonal patterns (Fekete et al., 1994; Leber and Sanes, 1995) (Figure 3D and S4E).

**Figure 3.**
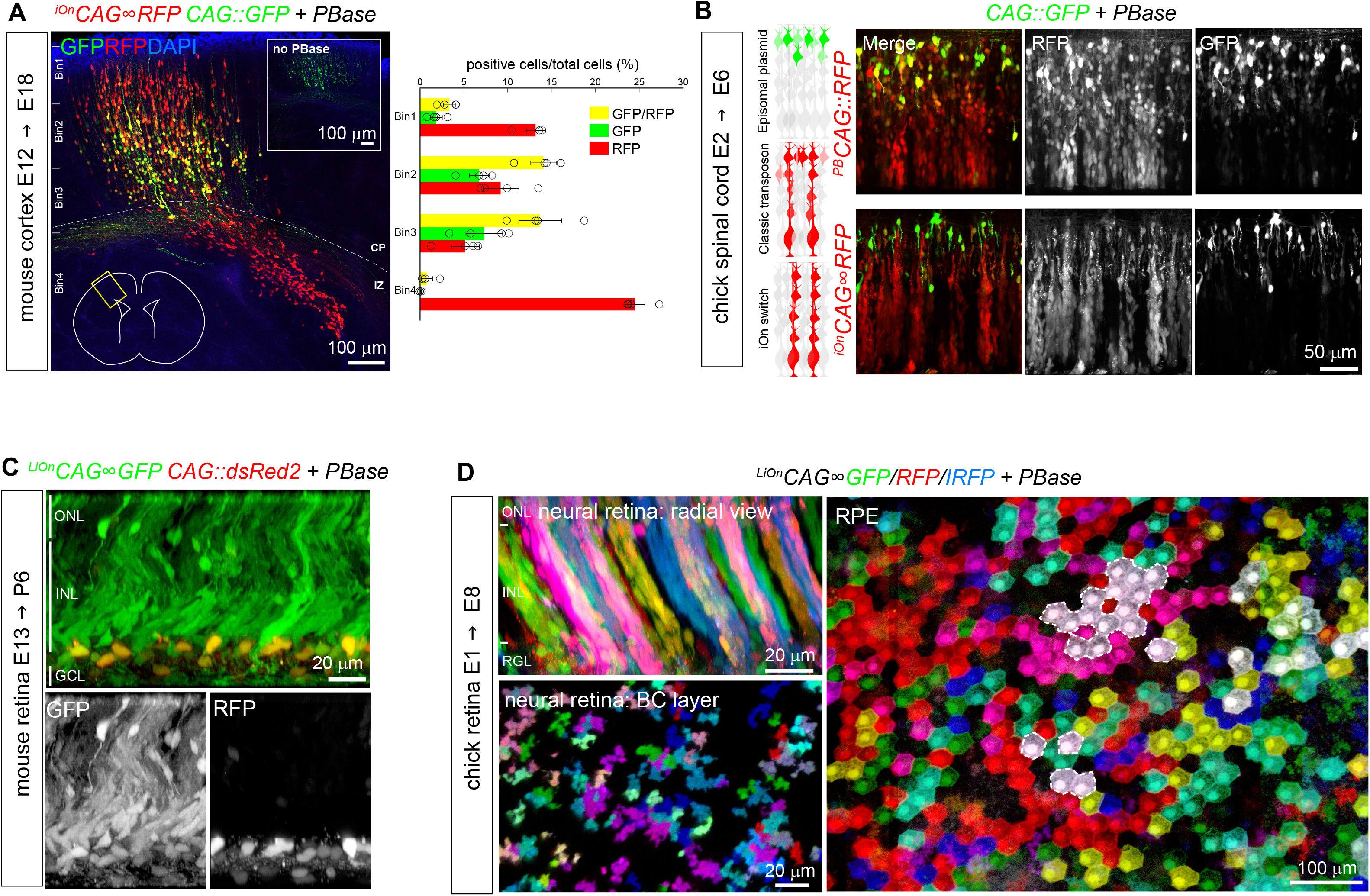
Cell lineage tracing and conditional expression by additive somatic transgenesis with iOn. (A) Fate mapping in the mouse cerebral cortex. Left: co-electroporation of an ^*iOn*^*CAG∞RFP* vector with PBase during neurogenesis (E12.5) yields streams of neurons migrating radially from the ventricular surface at E18.5, while an episome (*CAG∷GFP*) only marks those born shortly after electroporation. No red labeling is observed in absence of PBase (inset). CP, cortical plate. Right: quantification of labeled cells confirms that iOn-labeled cells (RFP, RFP/GFP) occupy all cortical layers, while most cells bearing episomal labeling settle in intermediate layers. (B) Longitudinal confocal views through E6 chick spinal cords electroporated at E2 with a classic transposon (top) or iOn vector (bottom) together with a control episome (*CAG∷GFP*). The iOn vector homogenously labels radially-migrating cells while the classic transposon also strongly labels isolated neurons, similar to the episomal vector. (C) Lineage tracing. ^*LiOn*^*CAG∞GFP* electroporation with PBase in the embryonic mouse retina during neurogenesis (E13) labels all retinal layers at P6, while expression from an episome (*CAG∷dsRed2*) only marks ganglion cells, born shortly after electroporation (ONL, INL, GCL: outer nuclear, inner nuclear and ganglion cell layers). (D) Multicolor clonal tracking. Radial view of a portion of neural retina (top) and en-face view of the bipolar cell layer (BC, bottom), and retinal pigmented epithelium (RPE, bottom) from an E8 chicken embryo electroporated with triple-color LiOn vectors at E1.5. FP combinations identify clones. See also Figures S4.

### Functional mosaic analysis with iOn

Beyond lineage tracing, the iOn strategy offers a mean to create sustained experimental perturbations of specific cell signaling pathways (Figure 4). To test this possibility, we assembled a LiOn vector co-expressing RFP and the intracellular domain of the Notch receptor (NICD), a well-known regulator of neural progenitor fate (Pierfelice et al., 2011) (Figure 4A). Electroporation of this plasmid in the embryonic chick spinal cord produced the effects expected for Notch pathway activation (Hammerle et al., 2011): marked reduction of neurogenesis and expansion of progenitors, compared to a ^*LiOn*^*CAG∞GFP* control vector (Figure 4A, B, Figure S5). Remarkably, these effects were still manifest 4 days after electroporation and were also augmented with respect to a transient perturbation. This experimental paradigm provided us with a unique opportunity to investigate the non-cell autonomous effects of an extended maintenance of neural stem cells (Figure 4C). To this aim, we generated color-coded mosaics by co-electroporating NICD-expressing (red) and control (green) iOn vectors, in which we measured the neurogenic output of GFP^+^/RFP^−^ (unperturbed) progenitors 4 days after the electroporation. Of note, such mosaic analysis cannot be conclusively undertaken with classic integrative vectors active in episomal form since cells unlabeled at the time of the analysis may have transiently expressed the transgene earlier on. Surprisingly, in the iOn mosaics, we observed that unperturbed spinal progenitors generated significantly expanded numbers of progeny (+314%, p< 0.005) compared to a control situation (Figure 4C). This reveals that neural stem cells of the embryonic spinal cord can modulate their output and compensate in a homeostatic manner for a reduction of neurogenic activity of their neighbors.

**Figure 4.**
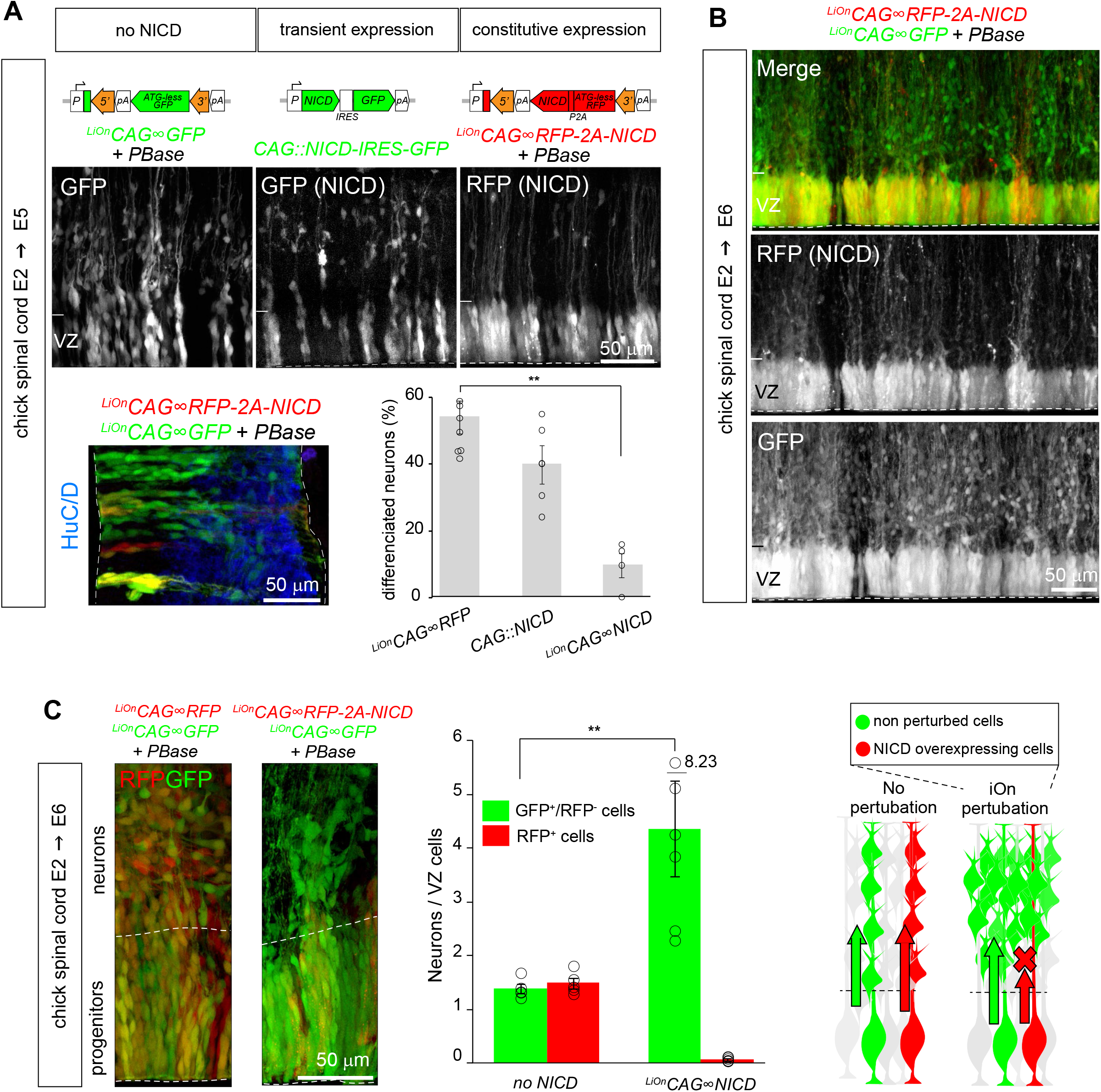
Functional mosaic analysis by somatic transgenesis and summary of iOn applications. (A) Top, longitudinal confocal views through E5 chick spinal cords 3 days after electroporation of a LiOn vector expressing the Notch intracellular domain (^*LiOn*^*CAG∞RFP-2A-NICD*), compared to transient NICD overexpression (*CAG∷NICD-IRES-GFP*) or control (^*LiOn*^*CAG∞GFP*). VZ: ventricular zone. Bottom, representative transverse section of E5 chick spinal cords electroporated with control (^*LiOn*^*CAG∞GFP*, green) and NICD-expressing LiOn vectors (^*LiOn*^*CAG∞RFP-2A-NICD*, red), immunostained for neuronal marker HuC/D (blue). Graph shows the percentage of HuC/D neurons among transfected cells with sustained vs. transient NICD expression (*CAG∷NICD*) and control (^*LiOn*^*CAG∞RFP*). Values and error bars show mean and s.e.m. from distinct embryos. A Kruskal-Wallis test indicated significant difference between control and ^*LiOn*^*CAG∞NICD* (p<0.01). (B) Longitudinal view through an E6 spinal cord co-electroporated with control (^*LiOn*^*CAG∞GFP*, green) and NICD-expressing LiOn vectors (^*LiOn*^*CAG∞RFP-2A-NICD,* red). Green cells migrate radially while most red cells remain at the ventricular surface (dotted line). (C) Non-cell autonomous effects of NICD expression. Left, E6 chick spinal cord transverse sections co-electroporated at E2 with a green control LiOn vector and a ^*LiOn*^*CAG∞RFP* (left) or ^*LiOn*^*CAG∞RFP-2A-NICD* plasmid (right). Middle, quantification of the ratio of GFP+/RFP- and RFP+ neurons and ventricular zone cells. Values and error bars represent mean and s.e.m. from distinct embryos (n≥5). A Mann-Whitney test indicates significant difference between control and ^*LiOn*^*CAG∞NICD* (p<0.005). Right, summary. Increased neurogenic output from green cells in NICD-perturbed condition reveals a homeostatic interactions among progenitors. See also Figures S5.

### Conditional control of transgene expression in somatic transfection experiments

Finally, iOn vectors are also of interest to assay and exploit transcriptional regulatory elements in a genomic configuration without the need to establish transgenic cell or animal lines. Cre/*lox* conditional labeling, widely applied in transgenic mice to target specific cell types, has so far been incompatible with somatic transfection approaches due to leakage from episomes. We devised a LiOn vector that reconstitutes an interrupted Cre recombinase gene upon PBase action (Figure 5A, left). Strikingly, this ^*LiOn*^*CMV∞Cre* plasmid presented no detectable leakiness in absence of transposase when transfected in cells expressing a *floxed* reporter transgene, but efficiently triggered Cre expression and recombination upon PBase action (Figure 5A, right). We applied this LiOn Cre/*lox* switch to trace the fate of a subset of retinal progenitor cells (RPCs) defined by expression of the Atoh7 transcription factor. At the population level, Atoh7^+^ RPCs are known to be biased towards the generation of ganglion cells for which they are essential (Wang et al., 2001), but also generate much larger numbers of photoreceptor, horizontal and amacrine cells (Brzezinski et al., 2012; Feng et al., 2010). How this is accounted for at the individual progenitor level is unclear. *Atoh7* regulatory sequences (Skowronska-Krawczyk et al., 2009), validated in separate experiments (Figure S6A), were incorporated in a LiOn Cre vector. Electroporation of this ^*LiOn*^*Atoh7∞Cre* vector at early stages of chick retinogenesis (E1.5) drove expression in progenies restricted to the outer, amacrine and ganglion layers of the retina at E8, while bipolar cells located in the inner nuclear layer were not labeled (Figure 5B). This demonstrated tight control of Cre production by the *Atoh7* element during retinal development and confirmed that chicken Atoh7^+^ progenitors are biased towards specific fates. To determine the potency of individual Atoh7^+^ progenitors, we then combined the ^*LiOn*^*Atoh7∞Cre* vector with stochastic multicolor lineage reporters encoded by Tol2 *Cytbow* and *Nucbow* transposons (Loulier et al., 2014) (Figure 5C). This approach marked groups of retinal cells organized as columns (Figure 5C, left), within which FP combinations further delineated clones whose cell type composition could be assigned based on layer position and morphology. Strikingly at E8, these Atoh7^+^-derived clones mostly comprised only a single ganglion cell, but frequently corresponded to pairs of nearby cells of a same type among photoreceptors and amacrine cells (Figure 5C, Figure S6B), thus indicating that individual Atoh7^+^ progenitors generate an important fraction of these two types of neurons (at least 43.6±4.44%, and 36.4±7.38%, respectively) through terminal symmetric division that result in identical fates.

**Figure 5.**
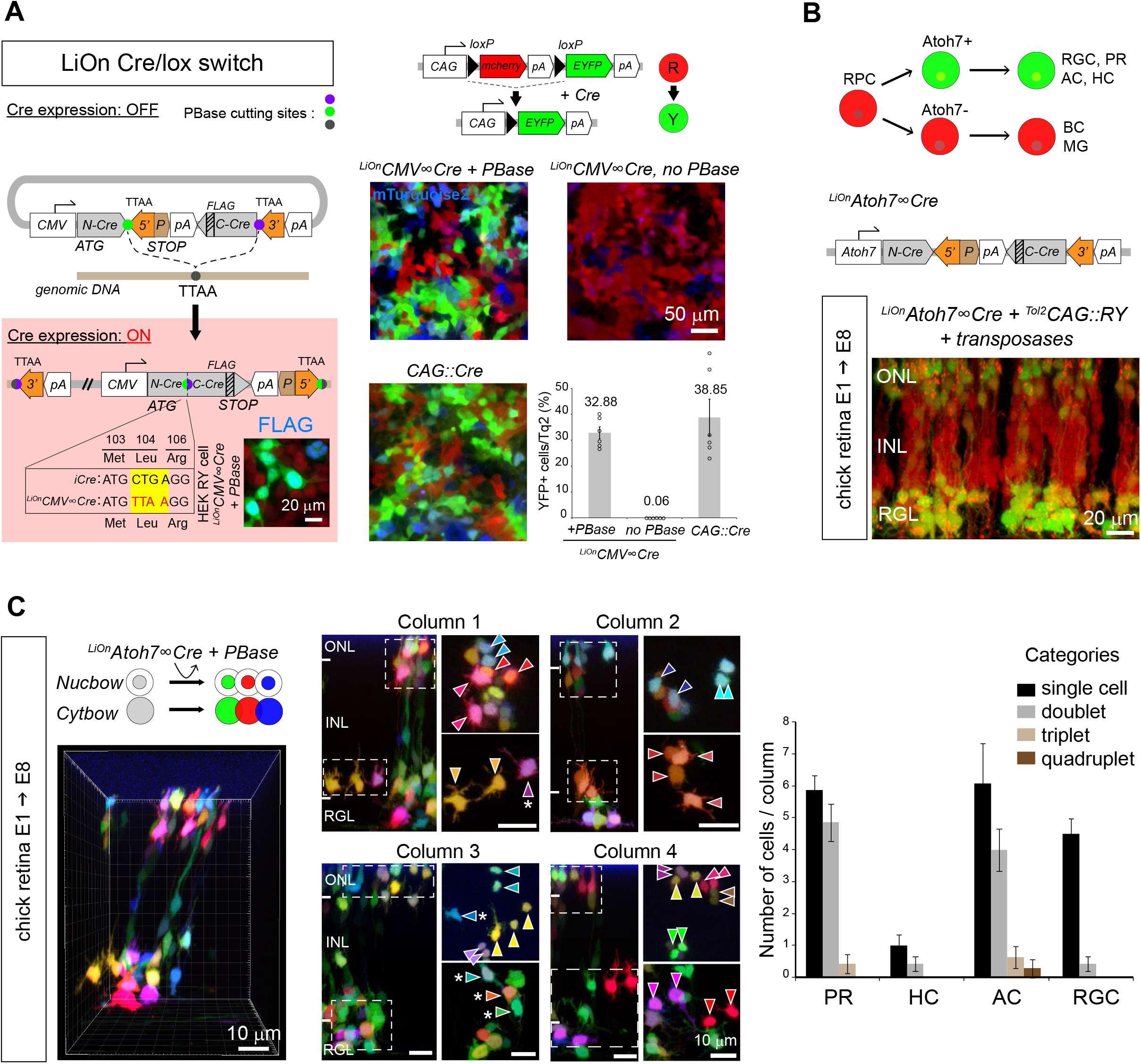
Intersectional Cre/lox recombination and analysis of the output of genetically identified neural progenitors with iOn. (A) Cre/*lox* conditional expression. Left, LiOn vector in which full translation of Cre, initially blocked, is activated by PBase; inset: Cre-FLAG immunodetection after transfection of ^*LiOn*^*CMV∞Cre* and PBase in HEK-RY cells stably expressing a *CAG∷RY* reporter switching from RFP to YFP expression under Cre action. Right, strict PBase-dependent recombination is observed 3 days after ^*LiOn*^*CMV∞Cre* transfection in HEK-RY cells (*CAG∷mTurquoise2:* transfection control). Graph shows mean and s.e.m. of replicates from 3 distinct experiments. (B) Radial view through an E8 chick retina co-electroporated at E1.5 with a Cre-expressing LiOn vector driven by *Atoh7* regulatory sequences (^*LiOn*^*Atoh7∞Cre*) and a ^*Tol2*^*CAG∷RY* transposon. Restricted recombination in the retinal ganglion (RGL) and outer nuclear (ONL) layers is observed. INL: inner nuclear layer. (C) Multicolor clonal analysis of Atoh7^+^ progenitor output. Optical sections and radial views of an E8 chick retina electroporated at E1.5 with ^*LiOn*^*Atoh7∞Cre* along with genome-integrating multicolor reporters (^*Tol2*^*CAG∷Nucbow and* ^*Tol2*^*CAG∷Cytbow*). Left: 3D view of a retinal column containing labeled neurons. Middle: Four retinal columns in which clonal pairs or sister cells of a same type can be identified based on expression of specific color marker combinations (arrowheads). Some cells not included in same-type pairs are also observed (asterisks). Column 1 corresponds to that shown in 3D in left panel. Right: quantification of the number of cells belonging to 1, 2, 3 or 4-cell clones within labeled photoreceptor (PR), horizontal (HC), amacrine (AC) and ganglion cells (RGC) in individual columns confirms a bias of Atoh7^+^ retinal progenitors that generate PRs and ACs towards terminal symmetric division patterns. Graph shows mean and s.e.m. of 14 columns reconstructed from 2 distinct embryos. See also Figures S6.

## DISCUSSION

We introduce a novel expression strategy that efficiently couples transgene activation to integration in the genome of host cells. iOn vectors rely on an unconventional arrangement of their elements to reconstitute a split transcriptional unit by DNA transposition. By canceling expression from non-integrated transgenes, the iOn switch solves two general problems associated with stable transfection using DNA vectors: the impossibility to identify transgenic cells directly after transfection due to episomes, and the adverse effects of episomal expression, including leakiness (Inoue et al., 2017), toxicity (Batard et al., 2001) and risks of genetic and epigenetic drift during selection (Liang and Zhang, 2013). In effect, iOn provides most of the advantages of retro- and lentiviral vectors without their cargo restriction, while offering the ease of use of DNA vectors and the capability of traceless excision of the piggyBac system (Fraser et al., 1996). Effective in all cell types and species tested here (human, mouse and chick), iOn can easily be adapted to different GOIs and promoters. For this purpose, maps of all plasmids built for this study, including a *CAG*-driven iOn vector equipped with a multicloning site, are presented in Table S1. The dependence of the iOn switch on transposition makes it ideally suited to report stable transgenesis in varied contexts (Figure 6). In cultured cells, the iOn switch essentially eliminates the need to select integration events through multiple rounds of cell division in stable transfection assays, making them as simple and rapid as transient approaches. As demonstrated here (Figures 1 and 2), its applications include rapid drug-free identification and sorting of stable integrants expressing one or multiple transgenes. In many uses, short term transfection experiments with iOn vectors will provide information equivalent to stable cell lines concerning gene expression, regulation and function. We anticipate that this will simplify and accelerate genetic screening procedures based on transposons (Kawakami et al., 2017). In vivo (Figures 3 and 4), transfection with iOn vectors can substitute for additive transgenesis in most if not all of its applications, with more accurate readout compared to previous somatic transgenesis approaches that can be polluted by expression from episomal vectors (Chen et al., 2014; Loulier et al., 2014). In particular, somatic transfection with iOn vectors provides a very efficient mean to track cell lineage in any organ accessible to electroporation, making them an ideal tool for studies in non-genetically tractable models. iOn also opens a simple route for functional mosaic analysis in vertebrate animals, normally requiring complex, time-consuming genetic manipulations (Pontes-Quero et al., 2017; Zong et al., 2005). In such experiments, the coupling of transgene integration and expression achieved with iOn ensures that marker expression in a cell reflects that of its entire lineage. This enabled us to create sustained mosaic perturbations of neurogenesis that reveal an intriguing homeostatic interplay among progenitors of the embryonic neural tube, capable of significantly influencing their output and reminiscent of that observed in some peripheral organs (Sharma et al., 2017; Stanger et al., 2007).

**Figure 6.**
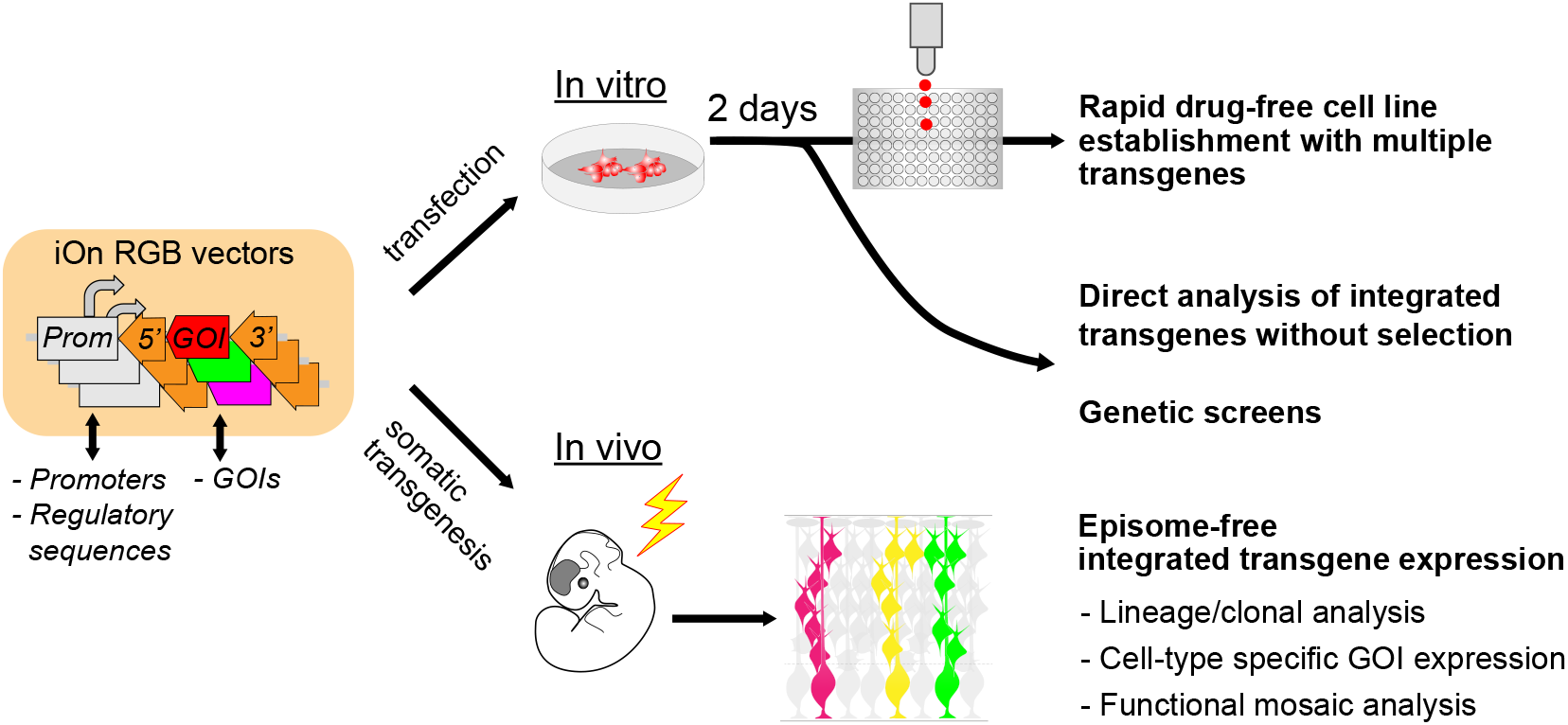
Summary of iOn potential applications. iOn vectors enable fast and reliable analysis of genome integrative events shortly after transfection both in vitro (top) and in vivo (bottom).

We also demonstrate that accurate cell-type specific Cre/*lox* recombination, an approach until now largely restricted to transgenic animal lines, can be achieved using electroporated iOn transgenes. This enabled us to trace in vivo the lineage of individual retinal progenitors defined by expression of the transcription factor Atoh7, revealing frequent terminal divisions that generate neurons of a same class, reminiscent of those observed with mouse cortical intermediate progenitors or Drosophila ganglion mother cells (Holguera and Desplan, 2018; Mihalas and Hevner, 2018). Such lineage pattern was particularly frequent for photoreceptors and amacrine cells, suggesting that progenitors restricted to generate these two cell types may exist in the chicken retina, in complement of those previously described for horizontal cells (Rompani and Cepko, 2008). Interestingly, terminal division patterns have also been observed with mouse retinal progenitors defined by expression of another bHLH transcription factor (Hafler et al., 2012), albeit in that case frequently generating distinct types of neurons. We hypothesize that, as observed in the mammalian cortex (Hevner, 2019), symmetric-fated terminal divisions observed in the chick retina could participate in establishing an appropriate balance among its different classes of neurons.

In conclusion, iOn represents a versatile expression strategy to report DNA vector integration that opens new avenues to engineer cells in cultured systems and directly probe the fate and regulation of progenitor cells in vivo, in the nervous system as well as in other organs accessible to transfection.

## ACKNOWLEDGEMENTS

We thank X. Morin and G. Orieux for scientific discussions, X. Nicol for the Kras sequence, J.-M. Matter for the *Atoh7* promoter, C. Marcelle for the NICD-expressing plasmid, and all members of the Livet lab for assistance. We thank L. Riancho and P.-H. Commère from the Saint-Antoine and Institut Pasteur cytometry facilities, A. Potey, C. Condroyer, S. Fouquet and Institut de la Vision core facilities. This work was funded by fellowships from the Uehara memorial and Naito foundations to T. K., the French ministry of research to F.M. and C.V., by the European Research Council (ERC-CoG n° 649117), and by Agence Nationale de la Recherche (contracts ANR-10-LABX-65, ANR-10-LABX-73-01 and ANR-15-CE13-0010-02). S.N. was funded by ATIP/Avenir and Association Française contre les Myopathies and A.R. by Genespoir.

## AUTHOR CONTRIBUTIONS

J.L., R.B. and T.K. conceived the iOn and LiOn switches. Initial experiments, R.B.; in vitro and chick spinal cord experiments, T.K.; retina experiments, F.M. and A.R.; cortical electroporation, T.K. and K.L.; iPS cell experiments, C.V. and S.N.; ES cell experiments, M.C.T. and S.V.-P.; cell culture and assistance with cloning, M. Le. and M. Lerat.; data analysis, T.K. and S.T.; manuscript redaction, T.K., S.T. and J.L. with input from all other authors.

## METHODS

### Cultured cells

Human embryonic kidney (HEK293T), HeLa and 3T3 cells were cultured in 10% fetal bovine serum in Dulbecco modified Eagle medium (DMEM, Life technologies).

Human induced pluripotent stem cells (iPS line WTSIi008-A, EBiSC, UK) were cultured in E8 medium (Life technologies) on Geltrex coating (Life technologies) and passaged with EDTA. Mouse ES cells (C57BL/6 x 129/Sv, line KH2) (Beard et al., 2006) were cultured on primary embryonic fibroblasts feeder cells.

### Mice

Swiss strain females (Janvier labs) were housed in a 12 hr light/12 hr dark cycle with free access to food, and animal procedures were carried out in accordance with institutional guidelines. Animal protocols were approved by the Charles Darwin animal experimentation ethical board (CEEACD/N°5). The date of the vaginal plug was recorded as embryonic day (E) 0.5 and the date of birth as postnatal day (P) 0.

### Chicken embryos

JA57 chicken fertilized eggs were provided by EARL Morizeau (8 rue du Moulin, 28190 Dangers, France) and incubated at 38 °C for the appropriate time in a FIEM incubator (Italy).

### DNA constructs

A schematized map of the plasmids designed for this study can be found in Table S1, along with restriction sites available to exchange GOIs and promoters. All iOn and control piggyBac vectors were assembled in a pUC57-mini plasmid backbone (Genscript Inc) using a combination of DNA synthesis (Genscript Inc), Gibson assembly (NEB) and standard restriction and ligation-based cloning. PCR for Gibson assembly was performed using CloneAmp HiFi PCR Premix (Clontech) and Q5 high-fidelity DNA polymerase (NEB). We used minimal piggyBac 5’ and 3’ TRs (Meir et al., 2011), with an additional 3 bp from the wild type transposon in the 3’ TR as in Loulier et al., 2014. We drove GOI expression with the strong eukaryotic *CAG* (Niwa et al., 1991) and *CMV* promoters as well as a 2145 bp fragment regulating expression of the chicken *Atoh7* gene (Skowronska-Krawczyk et al., 2009). GOIs were followed by a bovine growth hormone transcriptional terminator (pA1). In the final iOn vector design, a rabbit beta globin transcription terminator (pA2) was added upstream of the PB 3’TR to prevent cryptic episomal transcription. FPs used as GOI included RFP (mRFP1, Campbell et al., 2002), GFP (EGFP, Clontech) and IRFP (IRFP670, Shcherbakova and Verkhusha, 2013). In LiOn vectors, FP ORFs were split near the N terminus (Nt) in two opposite-oriented fragments that become reunited by transposition with incorporation of the TTAA footprint as indicated in Figure 2A. In the ^*LiOn*^*CMV∞Cre* vector, the Cre recombinase ORF was separated in Nt and Ct portions as in (Jullien, 2003), with incorporation of the TTAA footprint at a silent position (Leu104) and addition of a FLAG epitope (DYKDDDDK) at the Ct of the protein. To limit expression of the Cre Nt fragment prior to transposition, its coding sequence was positioned in frame (through the PB 5’ TR) with a PEST degron (Li et al., 1998) followed by a translational stop. The membrane-restricted GFP was generated by adding a short Kras tethering sequence (Averaimo et al., 2016) at the Ct end of EGFP using annealed oligonucleotides. To assay Cre activity, we designed a *floxed* reporter (^*Tol2*^*CAG∷loxP-mCherry-loxP-EYFP,* abbreviated as ^*Tol2*^*CAG∷RY*) in which expression switches from mCherry to EYFP upon recombination, framed with *Tol2* transposition endfeet to enable genomic integration. The ^*LiOn*^*CAG∞RFP-2A-NICD* vector was assembled by introducing a P2A cleavage sequence between the RFP and NICD ORFs to enable their co-expression. As non-integrative control vectors, we used a *CAG∷NICD-IRES-GFP* plasmid (Rios et al., 2011). Other plasmids used in this study included *CMV-*driven vectors expressing Cre, mTurquoise2 (Goedhart et al., 2012) and IRFP670 (Shcherbakova and Verkhusha, 2013) as well as *CAG*-driven vectors producing EGFP, mCerulean, dsRed2 (Clontech), the Tol2 transposase (Kawakami and Noda, 2004) and an optimized piggyBac transposase (hyPBase (Yusa et al., 2011).

### HEK293, HeLa and NIH 3T3 cell culture experiments

iOn and piggyBac plasmids were transfected in human HEK293, HeLa or mouse NIH 3T3 cells using cationic lipids. Except when otherwise noted, 1×10^5^ cells/well were plated in a 24-well dish and transfected at day 1 with 100 ng *iOn* vector with or without 20 ng of PBase-expressing plasmid (*CAG∷hyPBase)* using 0.7 μl of Lipofectamine 2000 reagent (Invitrogen). For triple-color labeling experiments, we used ng/well of each ^*LiOn*^*CAG∞FP* plasmid and 60 ng of PBase vector. To validate the ^*LiOn*^*CMV∞Cre* transgene, 50 ng of the corresponding plasmid was co-transfected with 10 ng of PBase vector in a HEK293 cell line stably expressing the ^*Tol2*^*CAG∷RY* reporter. This line was established by successive use of Tol2 transposition, drug selection with G418 (300 μg/ml, Sigma) and picking of RFP-positive clones. In some experiments, 50 ng of non-integrative plasmid expressing an FP marker distinct from the iOn vector (*CMV∷mTurquoise2*, *CMV∷IRFP* or *CAG∷GFP*) were applied as transfection control. For FACS analysis, transfections were performed in 6-cm dishes with scaled up concentrations. HEK293 cell viability after iOn plasmids transfection was assessed by dye exclusion with Trypan blue solution (0.4%, Sigma). FP expression was either assayed by flow cytometry, epifluorescence or confocal microscopy, or an Arrayscan high-content system (Thermo Fisher Scientific) (see below). For fixed observations, cells grown on 13 mm coverslips coated with collagen (50 μg/ml, Sigma) were immersed in 4% paraformaldehyde (PFA) in phosphate buffer saline (PBS) (Antigenfix, Diapath), rinsed in PBS and mounted in glycerol-based Vectashield mounting medium supplemented with DAPI (Vector labs). All images are representative of at least 3 independent experiments.

### FACS analysis and selection of HEK293 clones

For FACS analysis, HEK293 cells grown on 6-cm dishes were dissociated three days after transfection, stained with DAPI and analyzed on a MoFlo Astrios cell sorter (Beckman Coulter) using the following laser lines: 405 nm (DAPI), 488 nm (GFP), 561 nm (RFP), 640 nm (IRFP). 10000 cells were analyzed for each condition; non-fluorescent controls were prepared from mock-transfected cells stained with DAPI. For clonal experiments, HEK293 cells were sorted as single cells two days after transfection. Selection windows were chosen to include most of the FP-positive population and exclude negative cells. For 3-color cell sorting, we first selected live dissociated cells and subsequently selected RFP+, IRFP+ cells within the GFP+ population. Cells were then plated as single cells in 96-well plates and grown for 7-10 days in 200 μL of 10% FBS/DMEM medium mixed 1:1 with filtrated HEK293-conditioned medium. FP expression was assayed by epifluorescence microscopy or Arrayscan High-Content imaging (see below). Some positive clones were expanded in larger dishes for sequencing. To this aim, genomic DNA was isolated from a confluent 3.5- or 10-cm dish with the Nucleospin Tissue Kit (Macherey-Nagel). The rearranged region between the promoter and GOIs (500-600 bp) was amplified using CloneAmp HiFi PCR premix (Clontech) followed by Sanger sequencing (Genewiz, UK).

### Human iPS cell transfection and differentiation

For iOn labeling of differentiating iPS cells, colonies were dissociated with Accutase (Life Technologies) and replated in 96-well plates coated with poly-L-ornithine (20 μg/ml, Sigma P4957) and laminin (3 μg/ml, Sigma 23017-015). On day 2, cells were transfected with Dreamfect (OZBioscience) according to the manufacturer’s instructions. Cells were then differentiated as spinal motor neurons, fixed on day 14 with 4% PFA and stained with Tuj1 antibody as previously done (Maury et al., 2015). For iPS line generation, WTSIi008-A iPS cells were plated and transfected with Lipofectamine Stem Cell reagent (Invitrogen) according to the manufacturer’s protocol. Transfected cells were isolated by manual or EDTA passages, and homogeneous colonies were obtained 18 days after transfection (4 passages).

### Mouse ES cell transfection and clone selection

KH2 ES cells were transfected with ^*LiOn*^*CAG∞GFP-Kras* and *CAG∷hyPBase* plasmids (4 to 1 weight ratio) using Lipofectamine 2000 reagent. 48 hrs after transfection, GFP-positives cells (1.5%) were sorted using an Astrios MoFlo EQ cell sorter and plated at low density (10^3^ cells/ 10-cm dish) on feeder cells. After eight days, GFP-positives clones were picked under a fluorescent stereomicroscope (Zeiss Discovery V20).

### Mouse and chicken embryonic electroporation

*In utero* and *in ovo* electroporation in mouse and chicken embryos were performed as previously described (Loulier et al., 2014; Rebsam et al., 2009). A DNA mix containing 1-1.2 μg/μl of iOn vector, 0.5-1.2 μg/μl of non-integrative control plasmid and 0.2 μg/μl of *CAG∷hyPBase* plasmid supplemented with fast green dye was injected with a glass capillary pipette into one lateral ventricle or eye of E12.5 or E14.5 mice, or the optic cup or central spinal cord canal of E1.5 or E2 chick embryos, respectively. For multicolor labeling, the mix contained 1 μg/μl of each ^*LiOn*^*CAG∞FP* vector and 0.6 μg/μl of PBase vector. Embryos were left to develop until sacrifice. Tissues were fixed in 4% PFA. Postnatal mouse brains were sectioned at 200-μm thickness with a vibrating microtome (VT1000, Leica), while mouse retinas, chick E6 spinal cords and E6-E8 retinas were flat-mounted on glass slides. Samples were mounted in Vectashield medium and imaged with epifluorescence or confocal fluorescence microscopy.

### Immunostaining

For cell cultures: HEK293 cells plated on glass coverslips or iPS cells were fixed with 4% PFA, followed by washing in PBS and a 20-60 min blocking step at room temperature. Blocking solution for HEK293 and iPS cells respectively contained 10% normal goat serum (Sigma) or fetal bovine serum (Eurobio) and 0.5% or 0.2% Triton X-100 (Sigma). Cells were then incubated overnight at 4°C with primary antibody diluted in blocking solution (rabbit anti-FLAG, Sigma, 1:250 or mouse anti-Tuj1, Biolegend 1:500). After washing in PBS and incubation with secondary antibody (Alexa 647 anti-goat IgG 1:500, or Alexa 488 anti-goat IgG, 1:1000, Invitrogen) for 1 hr at room temperature, cells were washed again prior to mounting in Vectashield medium. For chicken spinal cords sections: embryos fixed for 1 hr in 4% PFA were equilibrated in 30% sucrose and embedded in TissueTek (Sakura), frozen on dry ice and stored at −80°C prior to cryostat sectioning (Microm HM560, 14 μm sections). After equilibration at room temperature, sections were washed in PBS before blocking in PBS-0.1% Triton-10% normal donkey serum (NDS) and overnight incubation with primary antibody (1:50, anti-HuC/D, Molecular Probes) in PBS-0.1% Triton-1% NDS. Following PBS washes, slides were incubated 1 hr with secondary antibody (Alexa 647 donkey anti-mouse, Invitrogen, 1:500) in the above buffer, washed and mounted with Vectashield medium.

### Fluorescence imaging and image analysis

Epifluorescence images were collected with a 10 × 0.6 NA or 20 × 0.7 NA objective on a Leica DM6000 microscope equipped with a VT1000 camera and separate filter cubes for GFP, RFP and IRFP. Confocal image stacks were acquired with 20 × 0.8 NA oil and 40 × 1.25 NA silicone objectives on an Olympus FV1000 microscope, using 440, 488, 515, 560, and 633 nm laser lines to excite CFP, GFP, YFP, RFP and IRFP/Alexa 647, respectively. For analysis with Arrayscan (Thermo Fisher Scientific), cells grown in 24- or 96-well plates were fixed 15 min with 4% PFA and stained with 300 nM DAPI prior imaging with the following laser lines: 386 nm (DAPI), 485 nm (GFP), 570 nm (RFP), and 650 nm (IRFP). Images of live ES cell clones were acquired using an EVOS FL auto inverted microscope (Life Technologies). Image analysis was performed with Fiji (Schindelin et al., 2012) and Imaris. Levels were uniformly adjusted across images with Adobe Photoshop.

### Statistical analysis

The number of samples analyzed is indicated in the figure legends. Statistical analyses were performed using R or GraphPad Prism software. Significance was assessed using χ^2^ (Figure 1F, 2C, S2C and S2E) and Kruskal-Wallis (one-way ANOVA on ranks) tests (Figure 4A and S6B), and non-parametric Mann-Whitney U test (Figure 4C). Data represent mean ± SEM, * p < 0.05; ** p < 0.01, *** p < 0.001.

### FACS analysis

10000 cells were analyzed for each condition; non-fluorescent controls were prepared from mock-transfected cells stained with DAPI.

---

**Figure S1.**
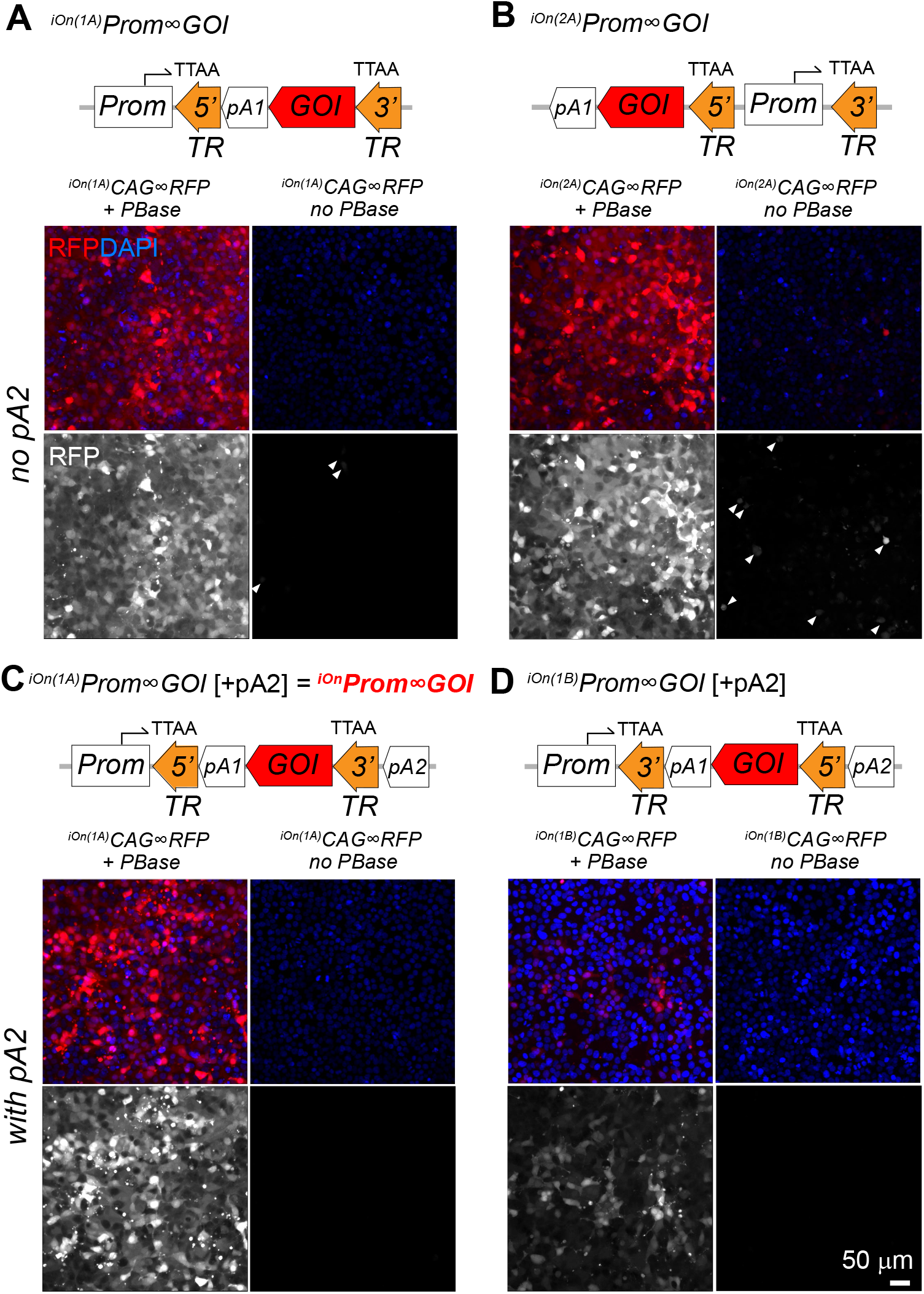
Design and test of transgene configurations for transposition-dependent GOI expression, related to Figure 1. (A, B) Top: two types of iOn transgene configurations (type 1, A and type 2, B) are possible depending on the positioning and relative orientation of the promoter and GOI. Bottom: test of the two configurations using *CAG* as promoter and *RFP* as GOI. 3 days after transfection in HEK293 cells, PBase-dependent expression is observed with both transgenes, with only weak leakiness in absence of PBase (arrowheads). (C, D) Top: two alternative designs of iOn switch (option A, C and option B, D) are possible depending on the positioning of the 5’ and 3’ transposon TRs. Schemes present the two alternatives on a type 1 configuration. A transcription terminator (pA2) is also positioned upstream of the GOI to reduce transcriptional leakiness prior to transposition. Bottom: test of the two configurations in HEK293 cells. PBase-dependent expression is observed with both transgenes, with reduced background expression compared to (A). The configuration presenting highest signal to background ratio shown in C (type 1Ap, C, hereafter denominated ^*PB*^*CAG∞RFP*), was selected for subsequent experiments.

**Figure S2.**
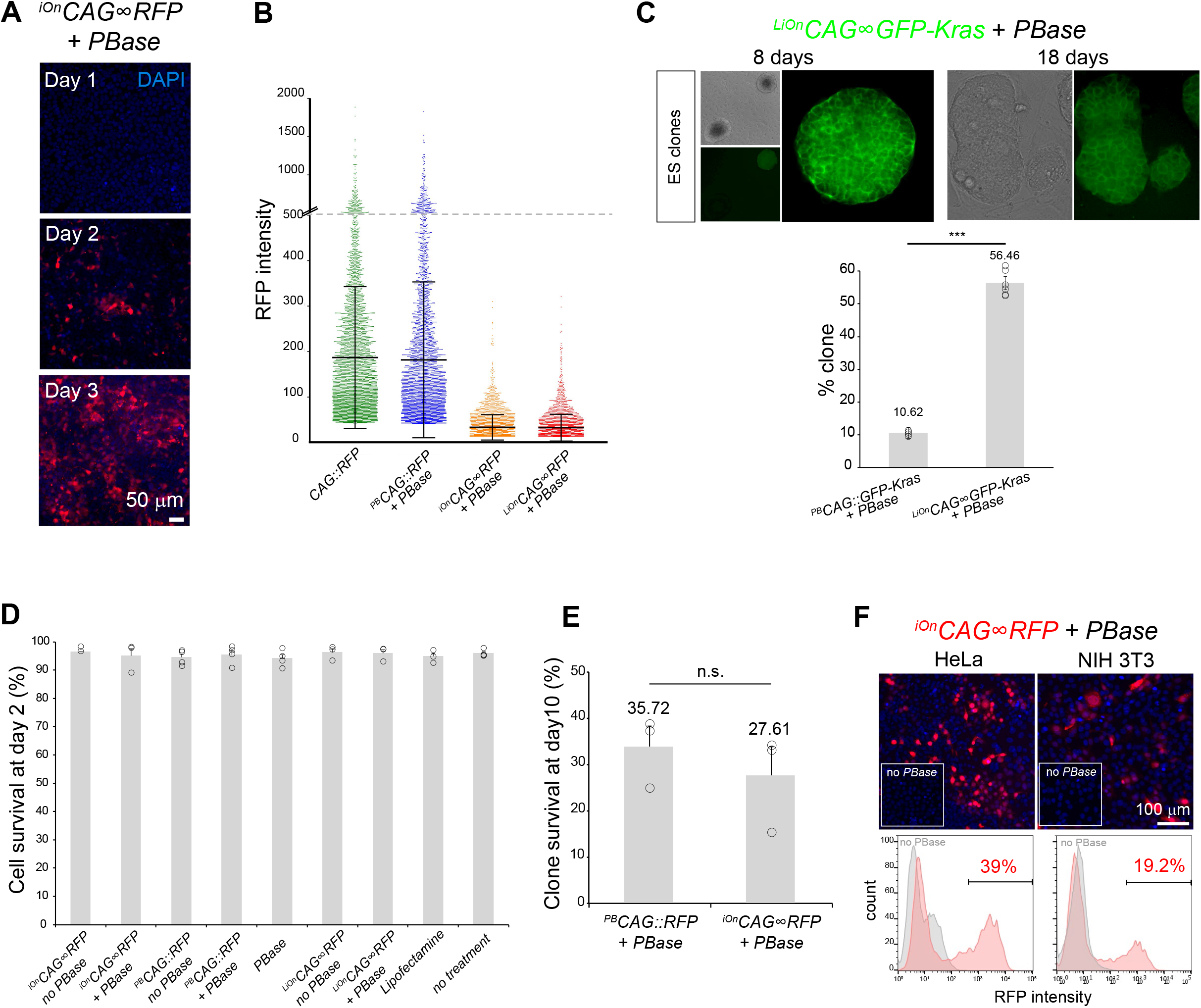
iOn vector expression in cultured cells, related to Figure 1. (A) Time-course of ^*iOn*^*CAG∞RFP* expression by epifluorescence imaging 1, 2, and 3 days after transfection in HEK293 cells in presence of PBase. (B) Comparison of expression levels from individual cells transfected with episomal (*CAG∷RFP*), classic transposon (^*iOn*^*CAG∞RFP* + *PBase*) and iOn vectors (^*iOn*^*CAG∞RFP* + *PBase* and ^*LiOn*^*CAG∞RFP* + *PBase*). Measures acquired 3 days after transfection on a high-content imaging platform. (C) Top: mouse ES cell clones grown for 8 and 18 days after sorting based on expression of a LiOn vector encoding a membrane-bound GFP (^*LiOn*^*CAG∞GFP-Kras* transfected 2 days prior to sorting). Left: low magnification picture showing GFP-positive and negative ES cell clones. Right: higher magnification of a positive clone showing membrane localization of GFP in all cells of the clone. Bottom: Compared to a classic transposon vector (^*PB*^*CAG∷GFP-Kras*), cells sorted based on ^*LiOn*^*CAG∞GFP-Kras* expression result in a higher yield of GFP-positive clones. Values and error bars represent the mean and s.e.m or four distinct replicates (dots). A χ2 test indicated a significant difference between the two situations (p<0.001). (D) Assessment of cell viability by Trypan blue assay 2 days after HEK293 cells transfection with iOn and control vectors. >95% survival is observed in all conditions. (E) Survival at 10 days of clones established from single RFP-positive cell sorted after transfection with ^*PB*^*CAG∷RFP* and ^*iOn*^*CAG∞RFP* in presence of PBase (measurements from the experiment presented in Fig. 1F). A χ2 test indicated non-significant difference between the two situations. (F) PBase-dependent expression from a ^*iOn*^*CAG∞RFP* vector in HeLa and NIH 3T3 cells 3 days after transfection in presence and absence of PBase (top: epifluorescence imaging, bottom: FACS analysis).

**Figure S3.**
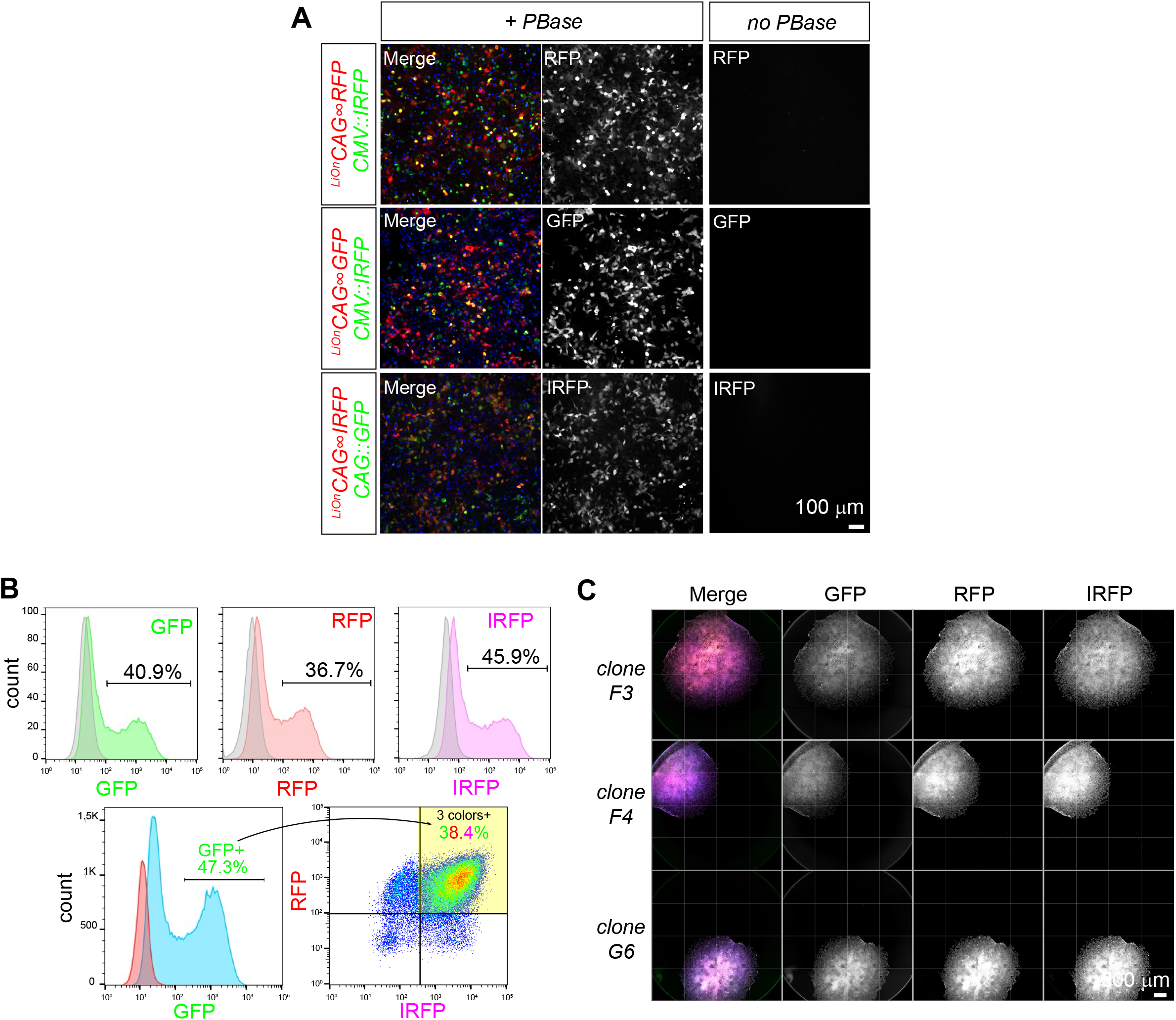
Multiplex transgenesis with iOn vectors, related to Figure 2. (A) Validation of three ^*LiOn*^*CAG∞FP* vectors respectively expressing RFP (top), EGFP (middle) and IRFP (bottom) in a PBase-dependent manner. HEK293 cells were imaged 3 days after transfection. Signal from LiOn vectors is encoded in red, while that of episomal vectors used as transfection control is shown in green (blue: DAPI staining). (B) Analysis and sorting of triple-labeled cells co-expressing the three ^*LiOn*^*CAG∞RFP* (GFP/RFP/IRFP) vectors. HEK293 cells were analy-zed 2 days after transfection. Top: FACS analysis of single channels (colored plots) from triple-transfected cells compared to cells transfected with a non-fluorescent plasmid (grey). Bottom: parameters for single cell sorting. Within the GFP positive population (47.3%, bottom left), RFP/IRFP double-positive cells (38.4%, bottom right) were selected and plated as single cells. (C) Examples of clones co-expressing the three ^*LiOn*^*CAG∞FP* vectors.

**Figure S4.**
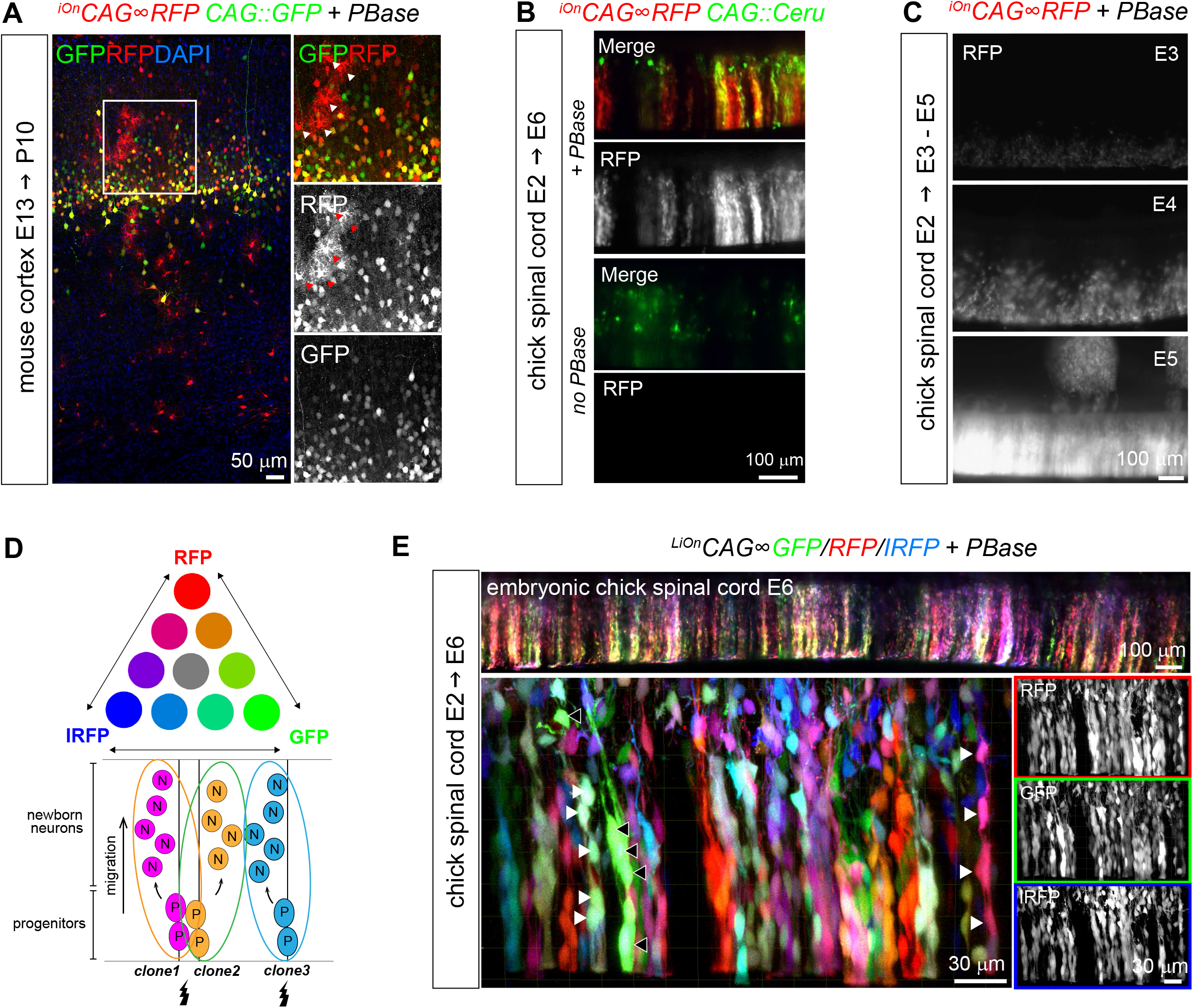
Application of iOn vectors in the vertebrate central nervous system, related to Figure 3. (A) At P10, astrocytes are also labeled with ^i*On*^*CAG∞RFP* but not with a *CAG∷GFP* plasmid electroporated at E13.5, demonstrating long term integration and expression of the iOn vector in neural progenitors and in their neuronal and glial descent (insets show enlarged view of boxed area, arrowheads point to protoplasmic astrocytes). (B) Validation of the iOn switch in vivo in the embryonic chicken spinal cord at E6. An episomal plasmid (*CAG∷Cerulean*, green) is found in isolated neurons born shortly after electroporation, while the iOn vector (red) labels radial clones of neurons migrating from the ventricular surface with the presence of PBase (top), but not the absence of PBase (bottom). (C) Time-course expression of ^*iOn*^*CAG∞RFP* with PBase by epifluorescence imaging 1, 2, and 3 days after electroporation in the embryonic chick spinal cord (whole mount views). (D) Top: Combinatorial expression of 3 FPs can generate several distinct color labels. Bottom: Expression of these labels in neurogenic progenitors can be used to resolve individual clones. (E) Wide-field epifluores-cence (top) and confocal views (bottom) from an E6 chick spinal cord electroporated at E2 with triple color LiOn mix (GFP/R-FP/IRFP) and PBase. FP combinations identify and resolve different clones of neural cells migrating radially from the ventricular surface (arrowheads).

**Figure S5.**
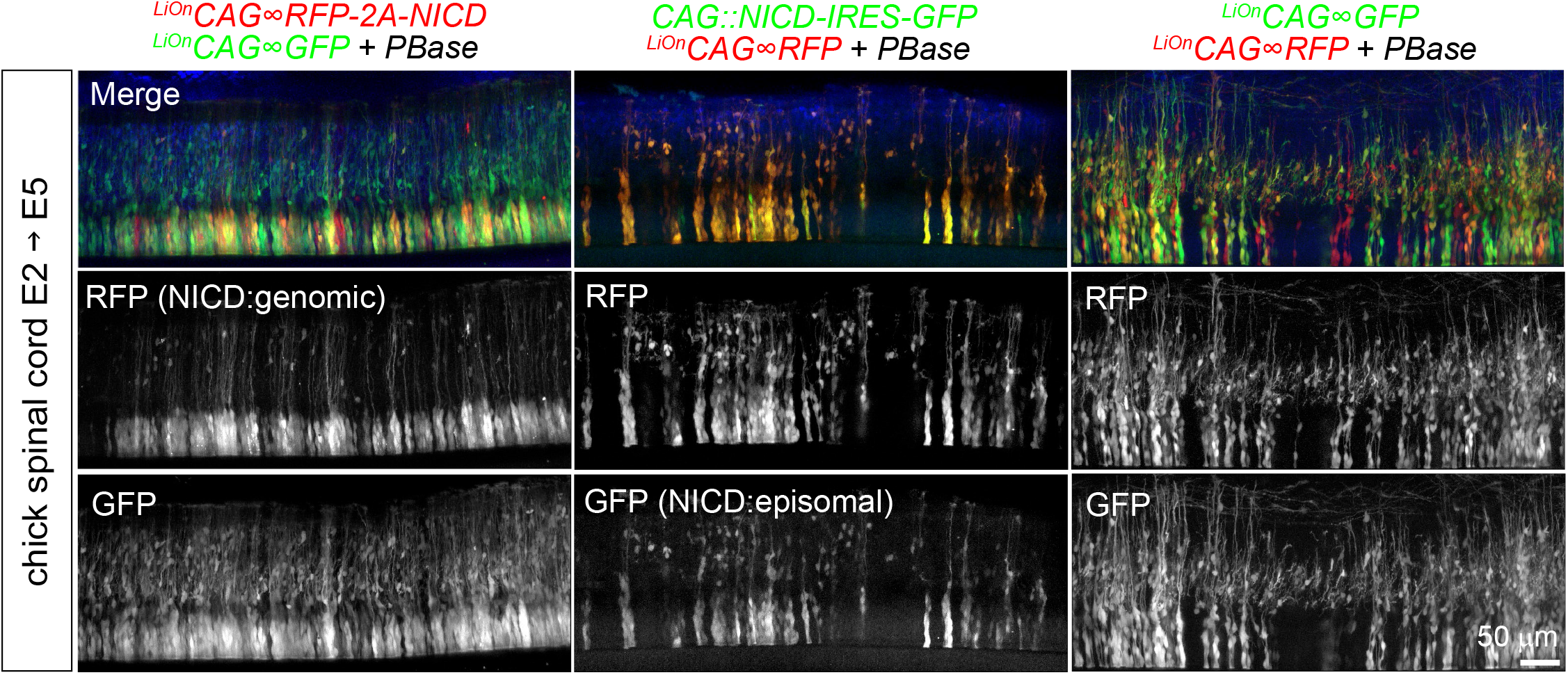
Long term perturbation of neural development by somatic transgenesis with iOn vectors, related to Figure 4. Comparison of the effects obtained in neural progenitors by expressing the Notch intracellular domain (NICD) from LiOn (left) and episomal vectors (middle) relative to a control situation right). Panels show longitudinal views through whole-mount embryonic chicken spinal cord preparations 3 days after electroporation, oriented such that the ventricular surface lines the bottom of the image. NICD-expressing plasmids were co-electroporated with non-perturbing plasmids as control. Permanent expression of NICD with LiOn (^*LiOn*^*CAG∞RFP-2A-NICD*, left) leads to an almost complete blockade of neuronal differentiation and radial migration (compare RFP signal with control ^*LiOn*^*CAG∞GFP* expression). Transient NICD expression (*CAG∷NICD-IRES-GFP*, middle) only results in partial effects compared with control (^*LiOn*^*CAG∞GFP/RFP*, right).

**Supplementary Figure 6 related with Figure 5.**
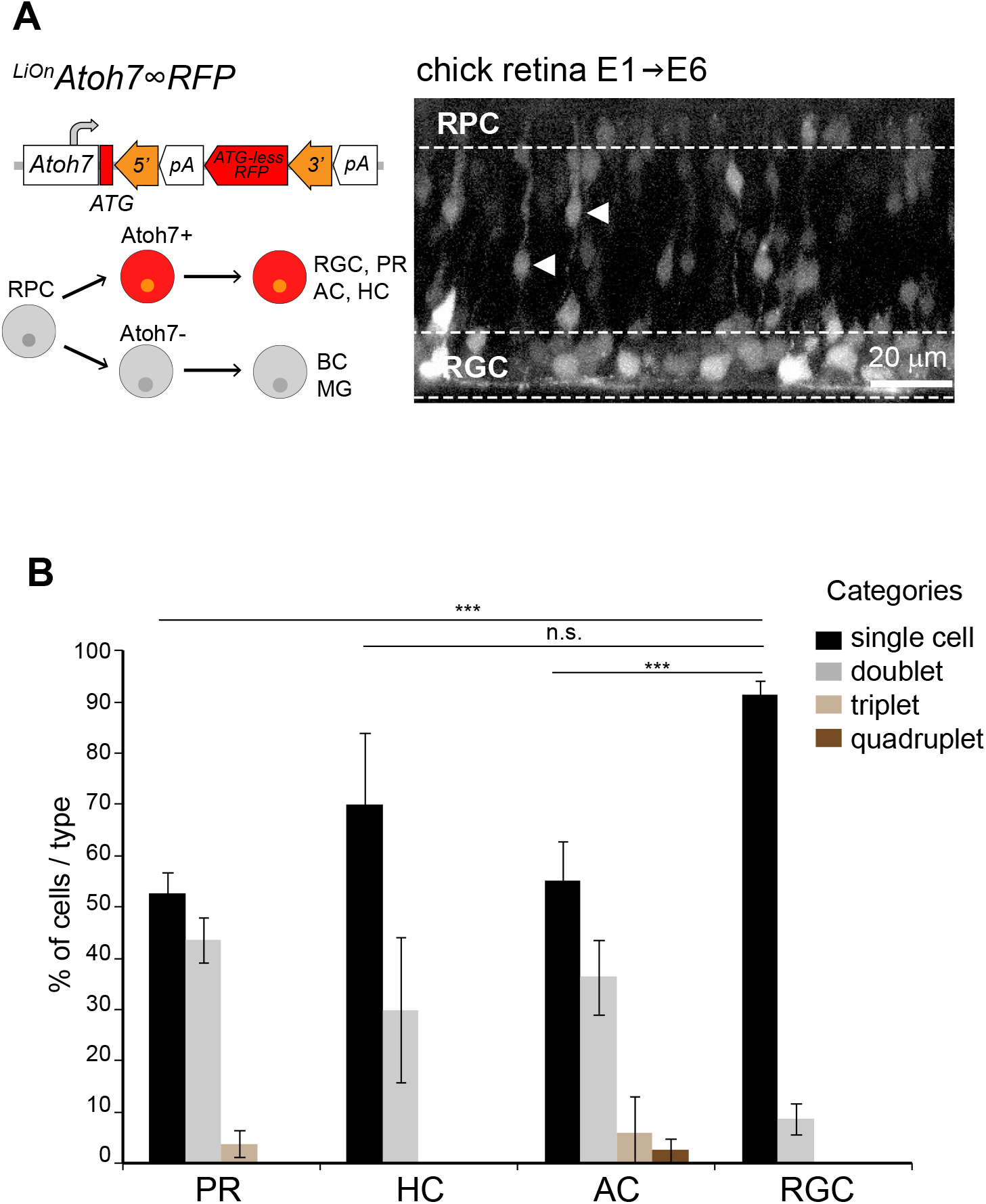
Long term perturbation of neural development by somatic transgenesis with iOn vectors. (A) View across the retina of an E6 chicken embryo, during the peak of RGC production and before the start of PR genesis, 5 days after electroporation of a ^*LiOn*^*Atoh7∞RFP* vector. Transgene expression is most prominent in the RGC (RGL) and outer nuclear (ONL) layers, while in intermediate layers only a fraction of labeled cells are visible, likely migrating towards the RGL at this intermediate stage (arrowheads). (B) Quantification of the percentage of Atoh7-derived retinal neurons belonging to 1, 2, 3 or 4-cell clones within labeled photoreceptor (PR), horizontal (HC), amacrine (AC) and ganglion cells (RGC) found in individual columns examined in Figure 5C. Graph shows mean and s.e.m. of 14 columns reconstructed from 2 distinct embryos. A Kruskal-Wallis test indicated significant difference between RGCs vs. PRs and HCs clonal category distributions (p<0.001).

**Table S1.** Schematic map of the transgenes designed for this study. All constructs were assembled in a pUC57-mini plasmid backbone. Restriction sites available to exchange GOIs and promoters are indicated. An iOn vector equipped with a multi-cloning site was also designed to facilitate the cloning of varied GOIs. pA1, pA2, pA3: bGH, rabbit beta-globin and SV40 transcription terminators. P: PEST degradation signal.

|  | Name | Scheme |
| --- | --- | --- |
| Test of iOn configurations | *iOn(1A)* CAG∞ RFP | |
| | *iOn(2A)* CAG∞ RFP | |
| | *iOn(1B)* CAG∞ RFP | |
| iOn (transcriptional switch) | *iOn* CAG∞ RFP | |
| | *iOn* CAG∞ MCS | |
| LiOn (translational switch) | *LiOn* CAG∞ GFP | |
| | *LiOn* CAG∞ RFP | |
| | *LiOn* CAG∞ IRFP | |
| Tagged LiOn | *LiOn* CAG∞ GFP-Kras | |
|  | Name | Scheme |
| Control PB transposons | *PB* CAG::GFP |  |
|  | *PB* CAG::RFP |  |
|  | *PB* CAG::IRFP |  |
| LiOn Cre | *LiOn* CMV∞ Cre | |
| NICD | *LiOn* CAG∞ RFP-2A-NICD | |
| Atoh7 promoter | *LiOn* Atoh7∞ Cre | |
| | *LiOn* Atoh7∞ RFP | |
|  | *Tol2*-based floxed reporter |  |

## REFERENCES

Akhtar, W., Jong, J. De, Pindyurin, A. V, Pagie, L., Meuleman, W., Ridder, J. De, Berns, A., Wessels, L.F.A., Lohuizen, M. Van, and Steensel, B. Van (2013). Resource Chromatin Position Effects Assayed by Thousands of Reporters Integrated in Parallel. Cell 154, 914–927.

Averaimo, S., Assali, A., Ros, O., Couvet, S., Zagar, Y., Genescu, I., Rebsam, A., and Nicol, X. (2016). A plasma membrane microdomain compartmentalizes ephrin-generated cAMP signals to prune developing retinal axon arbors. Nat. Commun. 7, 1–12.

Batard, P., Jordan, M., and Wurm, F. (2001). Transfer of high copy number plasmid into mammalian cells by calcium phosphate transfection. Gene 270, 61–68.

Beard, C., Hochedlinger, K., Plath, K., Wutz, A., and Jaenisch, R. (2006). Efficient method to generate single-copy transgenic mice by site-specific integration in embryonic stem cells. Genesis 44, 23–28.

Black, J.B., Perez-Pinera, P., and Gersbach, C.A. (2017). Mammalian Synthetic Biology: Engineering Biological Systems. Annu. Rev. Biomed. Eng. 19, 249–277.

Brzezinski, J.A., Prasov, L., and Glaser, T. (2012). Math5 defines the ganglion cell competence state in a subpopulation of retinal progenitor cells exiting the cell cycle. Dev. Biol. 365, 395–413.

Campbell, R.E., Tour, O., Palmer, A.E., Steinbach, P.A., Baird, G.S., Zacharias, D.A., and Tsien, R.Y. (2002). A monomeric red fluorescent protein. Proc Natl Acad Sci USA 99, 7877–7882.

Chen, F., and Loturco, J. (2012). A method for stable transgenesis of radial glia lineage in rat neocortex by piggyBac mediated transposition. J Neurosci Methods 207, 172–180.

Chen, F., Maher, B.J., and LoTurco, J.J. (2014). piggyBac transposon-mediated cellular transgenesis in mammalian forebrain by in utero electroporation. Cold Spring Harb. Protoc. 2014, 741–749.

Ding, S., Wu, X., Li, G., Han, M., Zhuang, Y., and Xu, T. (2005). Efficient transposition of the piggyBac (PB) transposon in mammalian cells and mice. Cell 122, 473–483.

Doench, J.G. (2018). Am I ready for CRISPR ? A user’s guide to genetic screens. Nat. Rev. Genet. 19, 67–80.

Ebrahimkhani, M.R., and Ebisuya, M. (2019). Synthetic developmental biology: build and control multicellular systems. Curr. Opin. Chem. Biol. 52, 9–15.

Fekete, D.M., Perez-Miguelsanz, J., Ryder, E.F., and Cepko, C.L. (1994). Clonal analysis in the chicken retina reveals tangential dispersion of clonally related cells. Dev Biol 166, 666–682.

Feng, L., Xie, Z., Ding, Q., Xie, X., Libby, R.T., and Gan, L. (2010). MATH5 controls the acquisition of multiple retinal cell fates. Mol. Brain 3, 1–16.

Fraser, M.J., Ciszczon, T., Elick, T., and Bauser, C. (1996). Precise excision of TTAA-specific lepidopteran transposons piggyBac (IFP2) and tagalong (TFP3) from the baculovirus genome in cell lines from two species of Lepidoptera. Insect Mol. Biol. 5, 141–151.

Goedhart, J., von Stetten, D., Noirclerc-Savoye, M., Lelimousin, M., Joosen, L., Hink, M.A., van Weeren, L., Gadella, T.W.J., and Royant, A. (2012). Structure-guided evolution of cyan fluorescent proteins towards a quantum yield of 93%. Nat. Commun. 3, 751.

Hafler, B.P., Surzenko, N., Beier, K.T., Punzo, C., Trimarchi, J.M., Kong, J.H., and Cepko, C.L. (2012). Transcription factor Olig2 defines subpopulations of retinal progenitor cells biased toward specific cell fates. Proc Natl Acad Sci USA 109, 7882–7887.

Hammerle, B., Ulin, E., Guimera, J., Becker, W., Guillemot, F., and Tejedor, F.J. (2011). Transient expression of Mnb/Dyrk1a couples cell cycle exit and differentiation of neuronal precursors by inducing p27KIP1 expression and suppressing NOTCH signaling. Development 138, 2543–2554.

Hevner, R.F. (2019). Intermediate progenitors and Tbr2 in cortical development. J. Anat.

Hirsch, T., Rothoeft, T., Teig, N., Bauer, J.W., Pellegrini, G., De Rosa, L., Scaglione, D., Reichelt, J., Klausegger, A., Kneisz, D., et al. (2017). Regeneration of the entire human epidermis using transgenic stem cells. Nature 551, 327–332.

Holguera, I., and Desplan, C. (2018). Neuronal specification in space and time. Science 362, 176–180.

Inoue, F., Kircher, M., Martin, B., Cooper, G.M., Witten, D.M., Mcmanus, M.T., Ahituv, N., and Shendure, J. (2017). A systematic comparison reveals substantial differences in chromosomal versus episomal encoding of enhancer activity. Genome Res. 27, 38–52.

Ivics, Z., Li, M.A., Mátés, L., Boeke, J.D., Nagy, A., Bradley, A., and Izsvák, Z. (2009). Transposon-mediated genome manipulation in vertebrates. Nat. Methods 6, 415–422.

Jullien, N. (2003). Regulation of Cre recombinase by ligand-induced complementation of inactive fragments. Nucleic Acids Res. 31, e131.

Kawakami, K., and Noda, T. (2004). Transposition of the Tol2 Element, an Ac-Like Element from the Japanese Medaka Fish Oryzias latipes, in Mouse Embryonic Stem Cells. Genetics 166, 895–899.

Kawakami, K., Largaespada, D.A., and Ivics, Z. (2017). Transposons As Tools for Functional Genomics in Vertebrate Models. Trends Genet. 33, 784–801.

Kondo, T., Imamura, K., Funayama, M., Tsukita, K., Miyake, M., Ohta, A., Woltjen, K., Nakagawa, M., Asada, T., Arai, T., et al. (2017). iPSC-Based Compound Screening and In Vitro Trials Identify a Synergistic Anti-amyloid beta Combination for Alzheimer’s Disease. Cell Rep. 21, 2304–2312.

Leber, S.M., and Sanes, J.R. (1995). Migratory paths of neurons and glia in the embryonic chick spinal cord. J Neurosci 15, 1236–1248.

Li, M.A., Turner, D.J., Ning, Z., Yusa, K., Liang, Q., Eckert, S., Rad, L., Fitzgerald, T.W., Craig, N.L., and Bradley, A. (2011). Mobilization of giant piggyBac transposons in the mouse genome. Nucleic Acids Res. 39, e148.

Li, X., Zhao, X., Fang, Y., Duong, T., Fan, C., Huang, C., Kain, S.R., and Jiang, X. (1998). Generation of Destabilized Green Fluorescent Protein as a Transcription Reporter Generation of Destabilized Green Fluorescent Protein as a Transcription Reporter. J. Biol. Chem. 273, 34970–34975.

Liang, G., and Zhang, Y. (2013). Perspective Genetic and Epigenetic Variations in iPSCs : Potential Causes and Implications for Application. Stem Cell 13, 149–159.

Loulier, K., Barry, R., Mahou, P., Le Franc, Y., Supatto, W., Matho, K.S.S.K.S.K.S.S., Ieng, S., Fouquet, S., Dupin, E., Benosman, R., et al. (2014). Multiplex Cell and Lineage Tracking with Combinatorial Labels. Neuron 81, 505–520.

Lu, I., Chen, C., Tung, C., Chen, H., Pan, J., Chang, C., Cheng, J., Chen, Y., Wang, C., Huang, C., et al. (2018). Identification of genes associated with cortical malformation using a transposon-mediated somatic mutagenesis screen in mice. Nat. Commun. 9, 2498.

Maury, Y., Côme, J., Piskorowski, R.A., Salah-Mohellibi, N., Chevaleyre, V., Peschanski, M., Martinat, C., and Nedelec, S. (2015). Combinatorial analysis of developmental cues efficiently converts human pluripotent stem cells into multiple neuronal subtypes. Nat. Biotechnol. 33, 89–96.

Meir, Y.J.J., Weirauch, M.T., Yang, H.S., Chung, P.C., Yu, R.K., and Wu, S.C.Y. (2011). Genome-wide target profiling of piggyBac and Tol2 in HEK 293: Pros and cons for gene discovery and gene therapy. BMC Biotechnol. 11, 28.

Merkle, F.T., Ghosh, S., Kamitaki, N., and Mitchell, J. (2017). Human pluripotent stem cells recurrently acquire and expand dominant negative P53 mutations. Nature 545, 229–233.

Mihalas, A.B., and Hevner, R.F. (2018). Clonal analysis reveals laminar fate multipotency and daughter cell apoptosis of mouse cortical intermediate progenitors. Development 145.

Mikuni, T., Nishiyama, J., Sun, Y., Kamasawa, N., and Yasuda, R. (2016). High-Throughput, High-Resolution Mapping of Protein Localization in Mammalian Brain by In Vivo Genome Editing. Cell 165, 1803–1817.

Nehme, R., Zuccaro, E., Ghosh, S.D., Zhanyan, F., Li, C., Sherwood, J.L., Pietilainen, O., Feng, G., and Eggan, K. (2018). Combining NGN2 Programming with Developmental Patterning Generates Human Excitatory Neurons with NMDAR-Mediated Synaptic Transmission Resource Combining NGN2 Programming with Developmental Patterning Generates Human Excitatory Neurons with NMDAR-Mediated. CellReports 23, 2509–2523.

Nishiyama, J., Mikuni, T., and Yasuda, R. (2017). Virus-Mediated Genome Editing via Homology-Directed Repair in Mitotic and Postmitotic Cells in Mammalian Brain. Neuron 96, 755–768.e5.

Niwa, H., Yamamura, K., and Miyazaki, J. (1991). Efficient selection for high-expression transfectants with a novel eukaryotic vector. Gene 108, 193–199.

Noctor, S.C., Flint, A.C., Weissman, T.A., Dammerman, R.S., and Kriegstein, A.R. (2001). Neurons derived from radial glial cells establish radial units in neocortex. Nature 409, 714–720.

Parr-Brownlie, L.C., Bosch-Bouju, C., Schoderboeck, L., Sizemore, R.J., Abraham, W.C., and Hughes, S.M. (2015). Lentiviral vectors as tools to understand central nervous system biology in mammalian model organisms. Front. Mol. Neurosci. 8, 14.

Pierfelice, T., Alberi, L., and Gaiano, N. (2011). Notch in the vertebrate nervous system: an old dog with new tricks. Neuron 69, 840–855.

Pontes-Quero, S., Heredia, L., Casquero-García, V., Fernández-Chacón, M., Luo, W., Hermoso, A., Bansal, M., Garcia-Gonzalez, I., Sanchez-Muñoz, M.S., Perea, J.R., et al. (2017). Dual ifgMosaic: A Versatile Method for Multispectral and Combinatorial Mosaic Gene-Function Analysis. Cell 170, 800––814.e18.

Price, J., Turner, D., and Cepko, C. (1987). Lineage analysis in the vertebrate nervous system by retrovirus-mediated gene transfer. Proc Natl Acad Sci U S A 84, 156–160.

Rebsam, A., Petros, T.J., and Mason, C.A. (2009). Switching retinogeniculate axon laterality leads to normal targeting but abnormal eye-specific segregation that is activity dependent. J. Neurosci. 29, 14855–14863.

Rios, A.C., Serralbo, O., Salgado, D., and Marcelle, C. (2011). Neural crest regulates myogenesis through the transient activation of NOTCH. Nature 473, 532–535.

Rompani, S.B., and Cepko, C.L. (2008). Retinal progenitor cells can produce restricted subsets of horizontal cells. Proc. Natl. Acad. Sci. U. S. A. 105, 192–197.

Schindelin, J., Arganda-Carreras, I., Frise, E., Kaynig, V., Longair, M., Pietzsch, T., Preibisch, S., Rueden, C., Saalfeld, S., Schmid, B., et al. (2012). Fiji: an open-source platform for biological-image analysis. Nat. Methods 9, 676–682.

Sharma, B., Ho, L., Ford, G.H., Quertermous, T., Reversade, B., Red-horse, K., Sharma, B., Ho, L., Ford, G.H., Chen, H.I., et al. (2017). Alternative Progenitor Cells Compensate to Rebuild the Coronary Vasculature in Elabela - and Apj - Deficient Hearts. Dev. Cell 42, 655–666.

Shcherbakova, D.M., and Verkhusha, V. V. (2013). Near-infrared fluorescent proteins for multicolor in vivo imaging. Nat. Methods 10, 751–754.

Skowronska-Krawczyk, D., Chiodini, F., Ebeling, M., Alliod, C., Kundzewicz, A., Castro, D., Ballivet, M., Guillemot, F., Matter-Sadzinski, L., and Matter, J.-M. (2009). Conserved regulatory sequences in Atoh7 mediate non-conserved regulatory responses in retina ontogenesis. Development 136, 3767–3777.

Stanger, B.Z., Tanaka, A.J., and Melton, D.A. (2007). Organ size is limited by the number of embryonic progenitor cells in the pancreas but not the liver. Nature 445, 886–891.

Suzuki, K., Tsunekawa, Y., Hernandez-Benitez, R., Wu, J., Zhu, J., Kim, E.J., Hatanaka, F., Yamamoto, M., Araoka, T., Li, Z., et al. (2016). In vivo genome editing via CRISPR/Cas9 mediated homology-independent targeted integration. Nature 540, 144–149.

Tewary, M., Shakiba, N., and Zandstra, P.W. (2018). Stem cell bioengineering: building from stem cell biology. Nat. Rev. Genet. 19, 595–614.

Wang, S.W., Kim, B.S., Ding, K., Wang, H., Sun, D., Johnson, R.L., Klein, W.H., and Gan, L. (2001). Requirement for math5 in the development of retinal ganglion cells. Genes Dev 15, 24–29.

Yang, N., Chanda, S., Marro, S., Ng, Y.-H.Y.-H., Janas, J.J.A., Haag, D., Ang, C.E., Tang, Y., Flores, Q., Mall, M., et al. (2017). Generation of pure GABAergic neurons by transcription factor programming. Nat. Methods 14, 621–628.

Yu, Y.C., He, S., Chen, S., Fu, Y., Brown, K.N., Yao, X.H., Ma, J., Gao, K.P., Sosinsky, G.E., Huang, K., et al. (2012). Preferential electrical coupling regulates neocortical lineage-dependent microcircuit assembly. Nature 486, 113–117.

Yusa, K., Zhou, L., Li, M.A., Bradley, A., and Craig, N.L. (2011). A hyperactive piggyBac transposase for mammalian applications. Proc. Natl. Acad. Sci. 108, 1531–1536.

Zong, H., Espinosa, J.S., Su, H.H., Muzumdar, M.D., and Luo, L. (2005). Mosaic analysis with double markers in mice. Cell 121, 479–492.

